# *Drosophila p53* isoforms have overlapping and distinct functions in germline genome integrity and oocyte quality control

**DOI:** 10.1101/2020.07.21.214692

**Authors:** Ananya Chakravarti, Heshani N. Thirimanne, Brian R. Calvi

## Abstract

p53 gene family members in humans and other organisms encode a large number of protein isoforms whose functions are largely undefined. Using *Drosophila* as a model, we find that a p53B isoform is expressed predominantly in the germline where it colocalizes with p53A into subnuclear bodies. It is only p53A, however, that mediates the apoptotic response to ionizing radiation in the germline and soma. In contrast, p53A and p53B both respond to meiotic DNA breaks and are required during oogenesis to prevent persistent germline DNA breaks, an activity that is more crucial when meiotic recombination is defective. We find that in oocytes with persistent DNA breaks p53A is required to activate a meiotic pachytene checkpoint. Our findings indicate that *Drosophila* p53 isoforms have DNA lesion and cell type-specific functions, with parallels to the functions of mammalian p53 family members in the genotoxic stress response and oocyte quality control.

## Introduction

The p53 protein is best known as a tumor suppressor that plays a central role in the response to DNA damage and other types of stress (Lane and Crawford 1979; Linzer and Levine 1979; Levine 2020). In response to stress, p53 mostly acts as a homotetrameric transcription factor to induce gene expression that elicits cell cycle arrest, apoptosis, or autophagy, although it also has other non-transcription factor activities (Levine 2020). It is now clear, however, that p53 regulates a growing list of other biological processes, including metabolism, stem cell division, immunity, and DNA repair (Levine 2019). Vertebrate genomes encode two other p53 paralogs, p63 and p73, which also have diverse functions in stress response and development (Jost et al. 1997; Yang et al. 1998; Dotsch et al. 2010; Candi et al. 2014). Adding to this complexity, each of these three p53 family encode a large number of isoforms which can form homo- or heterocomplexes, both within and among the different gene paralogs (Fujita et al. 2009; Aoubala et al. 2011; Joruiz and Bourdon 2016; Anbarasan and Bourdon 2019; Fujita 2019). However, the function of only a small subset of these isoform complexes have been defined. In this study, we use the *p53* gene in *Drosophila* as a simplified genetic system to examine the function of p53 isoforms and find that they have critical overlapping and distinct functions during oogenesis.

The *Drosophila melanogaster* genome has a single p53 family member (Ingaramo et al. 2018). Similar to human p53 (TP53), it has a C terminal oligomerization domain (OD), a central DNA binding domain (DBD) and an N terminal transcriptional activation domain (TAD), and functions as a tetrameric transcription factor (Jin et al. 2000; Ollmann et al. 2000). This single *p53* gene expresses four mRNAs that encode three different protein isoforms (Figure 1A) (Ingaramo et al. 2018). A 44 kD p53A protein isoform was the first to be identified and is the most well characterized (Brodsky et al. 2000; Jin et al. 2000). Later RNA-Seq and other approaches revealed that alternative promoter usage and RNA splicing results in a 56 kD p53B protein isoform, which differs from p53A by an 110 amino acid longer N-terminal TAD that is encoded by a unique p53B 5’ exon (Roy et al. 2010; Ingaramo et al. 2018) (Figure 1A). Because the p53A isoform differs from p53B by a shorter N terminus, p53A is also known as ΔNp53 (Dichtel-Danjoy et al. 2013). Another short p53E mRNA isoform is predicted to encode a protein of 38 kD that contains the DNA binding domain but lacks the longer N-terminal TADs of p53A and p53B (Roy et al. 2010; Zhang et al. 2015) (Figure 1A).

**Figure 1.**
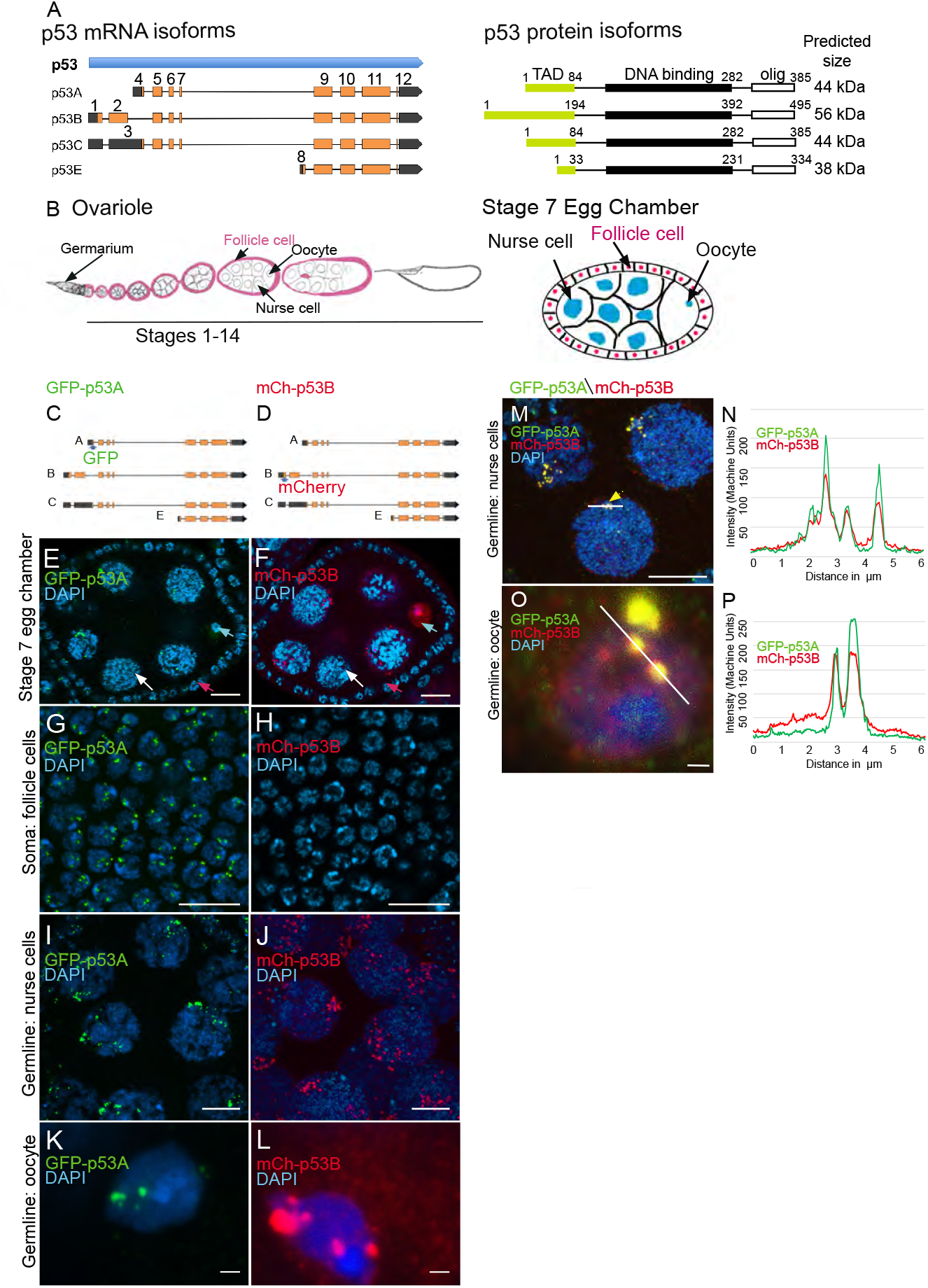
The p53B protein isoform is expressed in the germline where it colocalizes with p53A in nuclear bodies. (**A**) *Drosophila* p53 mRNA and protein isoforms. The left panel shows the four p53 mRNA isoforms with introns as lines, translated regions of exons as orange boxes, and 5’ and 3’ untranslated regions as black boxes. On the right are shown the p53 protein isoforms encoded by those four mRNA isoforms. Numbers indicate amino acid coordinates of transactivation domain (TAD) (green), DNA binding domain (black) and oligomerization domain (white). Note that p53A and p53C mRNAs encode the same protein. (**B**) Drosophila oogenesis: On the left is an illustration of one ovariole with egg chambers migrating posteriorly (to right) as they mature. The epithelial somatic follicle cells that surround each egg chamber are shown in pink. On the right is an enlarged longitudinal section through a stage 7 egg chamber showing somatic follicle cells surrounding germline nurse cells and oocyte. (**C, D**) Map of BAC transgenes with fluorescently-tagged p53 isoforms; GFP-p53A (**C**) and mCh-p53B (**D**). (**E-L**) Immunofluorescent detection of GFP-p53A (**E, G, I, K**) or mCh-p53B (**F, H, J, L**) expression in stage six egg chambers, with DNA counterstained with DAPI (blue). (**E, F**) A low magnification of a stage 6 egg chamber expressing GFP-p53A (**E**) and mCh-p53B. Subnuclear GFP-p53A bodies (**F**) were detected in germline nurse cells (white arrow), oocyte (blue arrow) and somatic follicle cells (red arrow), whereas mCh-p53B subnuclear bodies (**F**) were detected in all germline cells, but only rarely in somatic follicle cells (see Fig. S1). (**G-H**) Higher magnification images of p53A-GFP (**G, I, K**) and mCh-p53B (**H, J, L**) expression in somatic follicle cells (**G, H**), germline nurse cells (**I, J**), and oocyte (**K, L**). Scale bars are 10μm for **E-J**, 1μm for **K, L**. (**M-P**) GFP-p53A and mCh-p53B colocalize to germline subnuclear foci. Images of nurse cells (**M**) and oocyte (**O**) from a GFP-p53A / mCh-p53B female. The intensities in the red and green channel were quantified along the six micrometer lines shown in the images, with results shown for the indicated nurse cell nucleus (yellow arrow) (**N**) and oocyte (**P**). Scale bars are 10μm for **M**, and 1μm for **O**.

Like its human ortholog, *Drosophila p53* regulates apoptosis in response to genotoxic stress, but is now known to mediate other stress responses and developmental processes (Brodsky et al. 2000; Sogame et al. 2003; Wells et al. 2006; Dichtel-Danjoy et al. 2013; de la Cova et al. 2014; Napoletano et al. 2017; Tasnim and Kelleher 2018; Zhou 2019). To promote apoptosis, p53 protein directly induces transcription of several proapoptotic genes at one locus called H99 (Brodsky et al. 2000; Sogame et al. 2003; Zhou 2019). These early analyses of p53 function in apoptosis focused on the p53A isoform because the others had yet to be discovered. Using BAC rescue transgenes that were mutant for either p53A or p53B, we previously showed that in larval tissues it is the shorter p53A, and not p53B, that is both necessary and sufficient for the apoptotic response to DNA damage caused by ionizing radiation (Zhang et al. 2015). In contrast, when each isoform was overexpressed, p53B was much more potent than p53A at inducing proapoptotic gene transcription and the programmed cell death response, likely because of the longer p53B TAD (Dichtel-Danjoy et al. 2013; Zhang et al. 2015). Other evidence suggests that p53B may regulate tissue regeneration and has a redundant function with p53A to regulate autophagy in response to oxidative stress (Dichtel-Danjoy et al. 2013; Robin et al. 2019). It is largely unknown, however, why the Drosophila genome encodes a separate p53B isoform and what its array of functions are.

The p53 gene family is ancient with orthologs found in the genomes of multiple eukaryotes, including single-celled Choanozoans, which are thought to be the ancestors of multicellular animals (Rutkowski et al. 2010). Evidence suggests that the ancestral function of the p53 gene family was that of a p63-like protein in the germline, with later evolution of p53 tumor suppressor functions in the soma (Gebel et al. 2017; Levine 2020). In mammals, p63 mediates a meiotic pachytene checkpoint arrest in response to DNA damage or chromosome defects, and also induces apoptosis of a large number of oocytes with persistent defects, thereby enforcing an oocyte quality control (Di Giacomo et al. 2005; Suh et al. 2006; Gebel et al. 2017; Rinaldi et al. 2017; Rinaldi et al. 2020). It has been shown that in the *Drosophila* germline p53 regulates stem cell divisions, responds to programmed meiotic DNA breaks, and represses mobile elements (Lu et al. 2010; Wylie et al. 2014; Wylie et al. 2016). In this study, we have uncovered that the *Drosophila* p53A and p53B isoforms have redundant and distinct functions during oogenesis to protect genome integrity and mediate the meiotic pachytene checkpoint arrest, with parallels to the germline function of mammalian p53 family members in oocyte quality control.

## Results

### The p53B isoform is more highly expressed in the germline

Our previous results indicated that p53B does not mediate the apoptotic response to radiation in larval imaginal discs and brains (Zhang et al. 2015). One explanation for this lack of function was that p53B protein is expressed at very low levels in those somatic tissues (Zhang et al. 2015). Given the ancestral function of p53, we considered the possibility that p53B may be expressed and function in the germline. To address this question, we evaluated p53 isoform expression and function in the ovary. During *Drosophila* oogenesis, egg chambers migrate down a structure called the ovariole as they mature through 14 morphological stages (Figure 1B) (King 1970). Each egg chamber is composed of an oocyte and 15 sister germline nurse cells, all interconnected by intercellular bridges (Figure 1B) (Spradling 1993). The nurse cells become highly polyploid through repeated G / S endocycles during stages 1-10 of oogenesis, which facilitates their biosynthesis of large amounts of maternal RNA and protein that are deposited into the oocyte. The germline cells are surrounded by an epithelial sheet of somatic follicle cells that divide mitotically up until stage 6, and then undergo three endocycles from stages 7-10 (Calvi et al. 1998; Deng et al. 2001; Jia et al. 2015). Both germline and somatic follicle cell progenitors are continuously produced by germline and somatic stem cells that reside in a structure at the tip of the ovariole known as the germarium (King 1970; Drummond-Barbosa 2019).

To evaluate p53A and p53B expression during oogenesis, we used fly strains transformed with different p53 genomic BAC transgenes in which the p53 isoforms are tagged on their unique N-termini. In one strain, GFP is fused to p53A (GFP-p53A), while in another strain mCherry is fused to p53B (mCh-p53B), with each expressed under control of their normal regulatory regions in these genomic BACs (Figure 1C, D) (Zhang et al. 2015). To enhance detection, we immunolabeled these strains with antibodies that recognize GFP and mCherry. Immunofluorescent analysis of the somatic follicle cells revealed that GFP-p53A localized to nuclear bodies, ranging in size from ~0.25-1 μm, often in close proximity to the DAPI bright pericentric heterochromatin (Figure 1E,G). The expression of mCh-p53B, however, was only rarely detected in somatic follicle cells (< 1/ 50,000 cells) (Figure 1F,H, Figure S1). Thus, similar to our previous results in larval tissues, p53A is expressed at much higher levels than p53B in somatic cells (Zhang et al. 2015). In contrast, both GFP-p53A and mCh-p53B bodies were detected in all germline cells. Early stage nurse cells had one to a few p53 bodies, whereas later stage nurse cells had more bodies that were regionally distributed in the nucleus (Figure 1E, F, I, J). This dynamic pattern suggests that p53 bodies, like some other nuclear bodies, may associate with the polytene chromatin fibers that become dispersed in these nurse cells after stage 4 of oogenesis (Dej and Spradling 1999; Liu et al. 2006; White et al. 2007; Liu et al. 2009). The oocyte nucleus also had both GFP-p53A and Ch-p53B nuclear bodies, often appearing as one large (~1μm) and several smaller (~0.25μm) bodies (Figure 1K, L). In addition to distinct nuclear bodies, there were low levels of GFP-p53A and mCh-p53B dispersed throughout the nuclei of nurse cells and the oocyte.

We examined females with both GFP-p53A and mCh-p53B to address if they colocalize to the same nuclear bodies. In some cells co-expression of GFP-p53A reduced the expression of mCh-p53B, perhaps a manifestation of a protein trans-degradation effect that we had described previously (Zhang et al. 2014). Nonetheless, the results indicated that GFP-p53A and mCh-p53B colocalize to the same subnuclear bodies of both nurse cells and oocytes, although the ratio of these two isoforms differed somewhat among bodies (Fig 1M-P). mCh-p53B also co-localized with GFP-p53A in those rare follicle cells that expressed mCh-p53B (Figure S1B, C). Examination of the testis indicated that GFP-p53A and mCh-p53B are also expressed and localized to nuclear bodies in the male germline (Figure S2A-B’) (Mauri et al. 2008; Monk et al. 2012). Altogether, these results indicated that while the p53A isoform is expressed in both somatic and germline cells, the p53B isoform is primarily expressed in the germline.

### p53A is necessary and sufficient for the apoptotic response to ionizing radiation in somatic follicle cells

We next asked which of the p53 isoforms mediate the apoptotic response to DNA damage in the ovary. We had previously addressed this question in larval imaginal discs and brains using mutant BAC rescue transgenes (Zhang et al. 2015). For this study, we used CRISPR/Cas9 to create isoform-specific mutants at the endogenous *p53* locus. Since our previous study indicated that the short p53E is a repressor, we focused on making mutants of p53A and p53B isoforms to distinguish their different activities. The resulting *p53^A2.3^* allele is a 23bp deletion and 7bp insertion within the unique p53A 5’ exon (Figure 2A, Figure S3A). This deletion extends downstream into the first p53A intron removing both p53A coding sequence and first RNA splicing donor site (Figure 2A, Figure S3A) (Robin et al. 2019). This coding sequence and splice donor site are shared with p53C mRNA, which is predicted to encode a p53 protein isoform that is identical to that encoded by p53A mRNA (Figure 1A). Therefore, *p53^A2.3^* disrupts both p53A and p53C protein coding. The *p53^B41.5^* allele is a 14bp deletion plus 1bp insertion in the unique second coding exon of p53B, removing p53B coding sequence and creating a frameshift with a stop codon soon afterward (Figure 2B, Figure S3A). We had previously shown that p53A mRNA structure is perturbed and p53A protein is undetectable in homozygous *p53^A2.3^* animals, whereas p53B mRNA is still expressed (Figure S3B) (Robin et al. 2019). Conversely, RT-PCR indicated that in the *p53^B41.5^* strain the p53A isoform is still expressed (Figure S3B). Thus, *p53^A2.3^* and *p53^B41.5^* alleles are specific to each isoform and do not disrupt the function of the other isoform. This is in contrast to the *p53^5A-1-4^* null allele which deletes the common C-terminus of all the isoforms. To be clear about which of these isoforms are expressed from different *p53* alleles, we will annotate wild type *p53^+^* as (A+B+), the *p53^5A-1-4^* null allele as (A-B-), the p53A specific mutant *p53^A2.3^* as (A-B+) and the p53B specific mutant *p53^B41.5^* as (A+B-) (Figure 2A, B).

**Figure 2.**
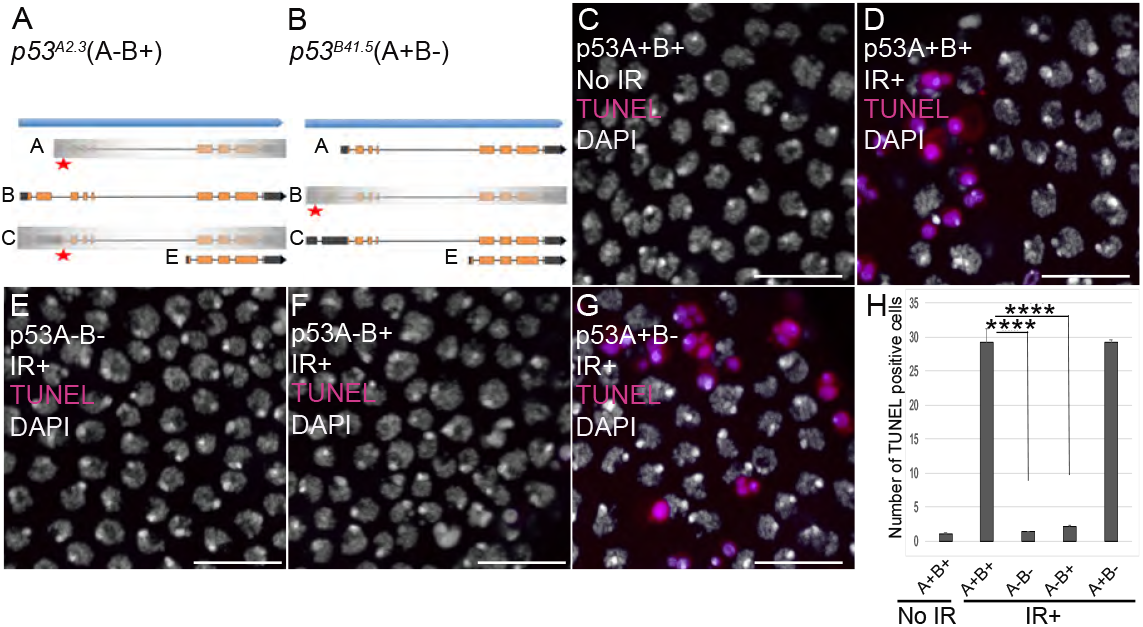
p53A is necessary and sufficient for IR-induced apoptosis in the soma. (**A-B**) Illustrations of the p53 isoform specific mutants created at the endogenous *p53* locus with CRISPR / Cas9. Each allele is a small deletion (red asterisk) in the unique 5’ coding exon of p53A (**A**) and p53B (**B**) mRNAs. The *p53^A2.3^* (A-B+) mutant impairs expression of isoforms p53A and p53C (gray shading) but not p53B, whereas the *p53^B41.5^* (A+B-) mutant eliminates expression of p53B (gray shading) but not p53A (see Fig. S3). (**C-H**) Apoptotic response to IR of stage 6 somatic follicle cells. Follicle cells were assayed for apoptosis by TUNEL (red) and DNA was stained with DAPI (gray). (**C, D**) TUNEL labeled follicle cells from a *p53^+^* (A+B+) wild type female without (**C**) or four hours after IR (**D**). (**E-G**) TUNEL labeling of follicle cells after IR from *p53^5A-1-4^* (A-B-) null (**E**), *p53^A2.3^* (A-B+) (**F**), and *p53^B41.5^* (A+B-) (**G**) mutant females. Scale bars are 10μm. (**H**) Quantification of the average number of TUNEL labeled follicle cells in stage 6 egg chambers for the genotypes and treatments shown in **C-G**. Averages are based on 10 egg chambers per genotype with two biological replicates. Error bars are S.E.M. ****: p<0.0001 by unpaired Student’s t test.

To determine which *p53* isoforms mediate the apoptotic response to DNA damage, we irradiated adult females from these strains with 40 Gray (Gy) of ionizing radiation (IR) and evaluated cell death four hours later by TUNEL. We focused on the follicle cells in the mitotic cycle up until stage 6 because we had previously shown that endocycling follicle cells in later stage egg chambers repress the p53 apoptotic response to DNA damage (Mehrotra et al. 2008; Hassel et al. 2014; Zhang et al. 2014; Qi and Calvi 2016). In wildtype *p53^+^* (A+B+) ovaries that express both isoforms, approximately 30% of mitotic cycling follicle cells were TUNEL positive, whereas the *p53^A2.3^* (A-B+) mutant had very few TUNEL-positive follicle cells (~1%), which was not significantly different than irradiated *p53^5A-1-4^* (A-B-) null or unirradiated controls (Figure 2C-F, H). In contrast, the *p53^B41.5^* (A+B-) mutant strain had 30% TUNEL-positive follicle cells, a fraction similar to that of wild type (Figure 2D, G, H). These results suggested that p53A, but not p53B, is required for the apoptotic response to DNA damage. A possible caveat, however, is that both the *p53^A2.3^* and *p53^B41.5^* alleles also delete part of a non-coding RNA of unknown function (CR46089), which overlaps the 5’ end of *p53* and is transcribed in the opposite direction (Roy et al. 2010; Thurmond et al. 2019). Given that this noncoding RNA is disrupted in both alleles, its disruption cannot explain the impaired apoptosis specifically in the *p53^A2.3^* allele. Moreover, similar results were obtained when *p53^A2.3^* or *p53^B41.5^* alleles were transheterozygous to the *p53^5A-1-4^* null allele that does not delete portions of this non-coding RNA. These results strongly suggest that mutation of non-coding RNA CR46089, or possible cryptic mutations on the *p53^A2.3^* and *p53^B41.5^* chromosomes, are not contributing to the apoptotic phenotypes. Thus, the p53A protein isoform is both necessary and sufficient for the apoptotic response to IR in somatic ovarian follicle cells.

### p53A is necessary and sufficient for the apoptotic response to ionizing radiation in the female germline

The low level of expression of p53B in somatic tissues may explain why it does not mediate the apoptotic response. We wondered, therefore, whether p53B participates in the apoptotic response in the germline where it is more highly expressed. Given that endocycling nurse cells and the meiotic oocyte repress p53 apoptosis, we analyzed the apoptotic response of mitotically-dividing germline cells during early oogenesis in the germarium (Mehrotra et al. 2008; Hassel et al. 2014; Zhang et al. 2014; Qi and Calvi 2016). At the anterior tip of the germarium, the germline stem cells (GSCs) reside in a niche and divide asymmetrically into a GSC and cystoblast (CB) (Figure 3A) (Hinnant et al. 2020). This cystoblast and its daughter cells undergo four rounds of divisions with incomplete cytokinesis as they migrate posteriorly through germarium region 1, finally resulting in an interconnected 16-cell germline cyst. (Figure 3A) (Drummond-Barbosa 2019; Hinnant et al. 2020). In region 2a, multiple cells in the cyst initiate meiotic breaks and synaptonemal complex formation, but only one cell is eventually specified to be the oocyte, with the 15 other cells of the cyst destined to become nurse cells that enter a polyploid endocycle by germarium region 3 (stage 1 of oogenesis) (Figure 3A). To evaluate which cells in the germarium express GFP-p53A and mCh-p53B, we co-labeled with an antibody against the fly adducin protein ortholog called Hu-li tai shao (Hts), which labels a spherical cytoplasmic spectrosome in GSCs, and a cytoskeletal structure called the fusome that branches through the ring canals that connect the 16 cells of a germline cyst (Figure 3A) (Lin et al. 1994). Similar to later stages of oogenesis, both GFP-p53A and mCh-p53B were expressed in GSCs and their daughter germline cells of the germarium where the p53 isoforms colocalized in distinct p53 nuclear bodies (Figure 3B, C).

**Figure 3.**
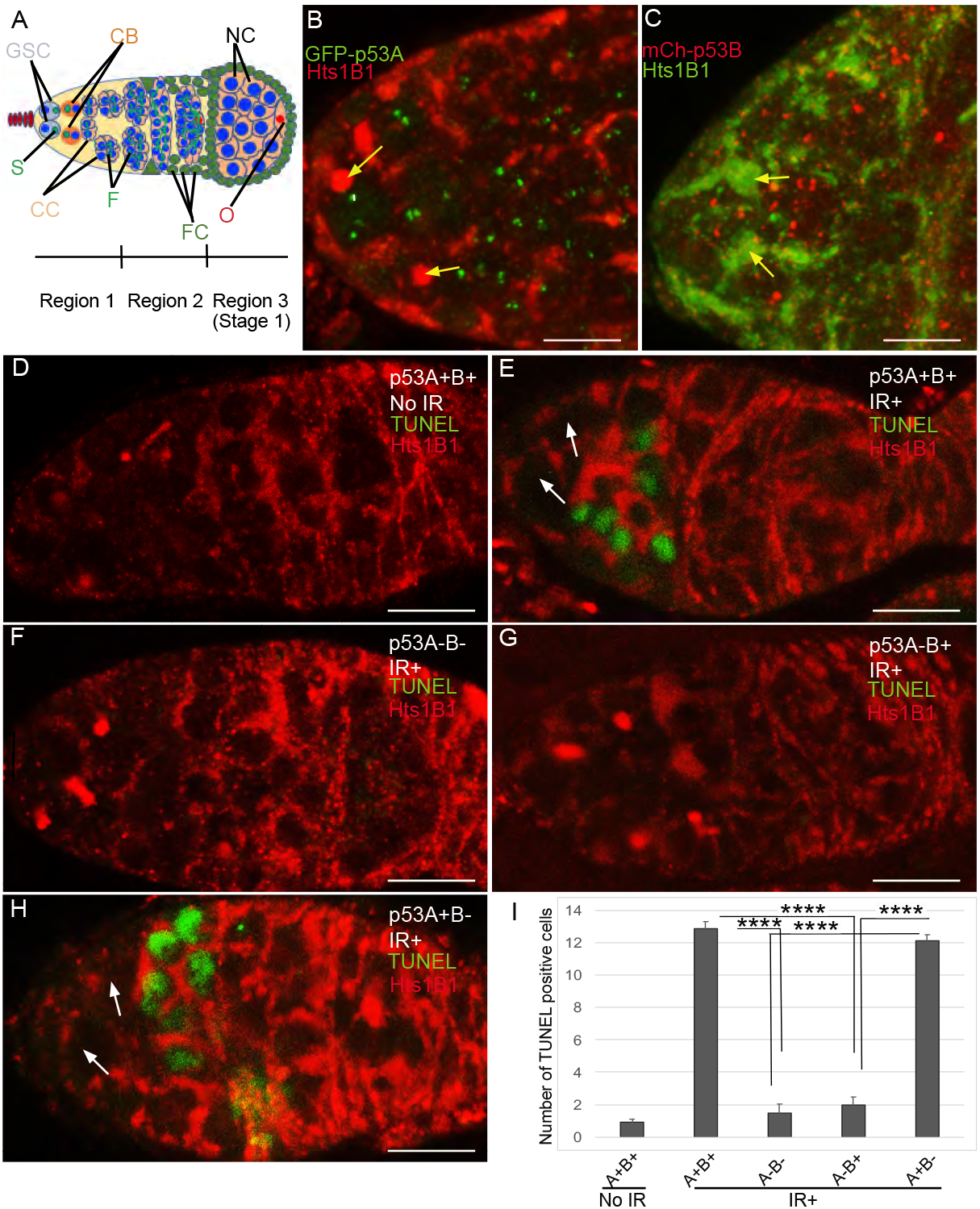
p53A and p53B are expressed in the early female germline, but only p53A is required for IR-induced germline apoptosis. (**A**) Illustration of the germarium. Region 1 of the germarium contains the female germline stem cells (GSC), their primary daughter cystoblasts (CB), and subsequent dividing cystocytes (CC). The GSCs contain a spherical cytoskeletal spectrosome (S) whereas the interconnected cystocytes (CC) contain a branched fusome (F, green lines). In region 1 / early region 2, the cystocytes undergo four incomplete divisions resulting in a 16 cell cyst interconnected by ring canals. In region 2, these cells begin meiosis including the initiation of DNA breaks. Thereafter, one cell is specified to become the oocyte (O) and proceed through meiosis to prometaphase of MI, whereas the other fifteen cells adopt the nurse cell (NC) fate and begin to endocycle. The germline cyst is surrounded by somatic follicle cells (FC, green) to form an egg chamber in germarium region 3 (oogenesis stage 1). (**B, C**) Expression of GFP-p53A (**B**) and mCh-p53B (**C**) in subnuclear bodies of GSC and cystocytes of the germarium. The GSCs were identified by the presence of the spectrosome (yellow arrows). Scale bars are 5μm (**D-E**) TUNEL (green) and anti-Hts labeling (red) in germaria from *p53^+^* (A+B+) wild type females without IR (**D**) or with IR (**E**). Spectrosomes and fusomes were labeled with anti-Hts antibody to identify GSC and cystocytes. (**F-H**) TUNEL after IR of *p53^5A-1-4^* (A-B-) null (**F**), *p53^A2.3^* (A-B+) (**G**) and *p53^B41.5^* (A+B-) (**H**) females. The GSCs (white arrows in E and H) were not TUNEL positive. Scale bars are 10μm (**I**) Quantification of the average number of TUNEL labeled cystocytes in region 1 of the germarium for the genotypes and treatments shown in **D-H**. Averages are based on 10 germaria per genotype and three biological replicates. Error bars are S.E.M. ****: p<0.0001 by unpaired Student’s t test.

To determine which p53 isoforms are required for IR-induced germline apoptosis, we irradiated wild type and *p53* mutant females with 40 Gy of gamma rays and TUNEL labeled their ovaries four hours later. In wild type *p53^+^* (A+B+) controls, there were an average of ~13 TUNEL-positive germline cells in region 1 of each germarium (Figure 3D,E,I). Although earlier GSCs and later meiotic cells express both p53 isoforms, they did not label with TUNEL (Figure 3E). In *p53^5A-1-4^* (A-B-) null ovaries, only ~1 germline cell per germarium was TUNEL-positive in region 1, a number similar to that in unirradiated controls, indicating that most of the germline cell death four hours after IR is p53-dependent (Figure 3 D, F, I). Similar to *p53^5A-1-4^* null, the *p53^A2.3^* (A-B+) mutant also had ~1 TUNEL-positive germline cell per germarium (Figure 3G, I). In contrast, the *p53^B41.5^* (A+B-) mutant ovaries had ~12 TUNEL positive cells / germarium, a number similar to that in wild type and significantly greater than that in *p53* null and *p53^A2.3^* mutants (Figure 3H, I). These results suggest that, despite p53B expression, it is the p53A isoform that is necessary and sufficient for the apoptotic response to IR in the germline.

To further evaluate p53 isoform function, we determined whether p53A or p53B protein isoforms induce transcription of proapoptotic genes after IR. Previous studies showed that among the proapoptotic p53 target genes, the gene *hid* plays a prominent role for inducing germline apoptosis in response to DNA damage (Xing et al. 2015; Park et al. 2019). We therefore used a GFP promoter-reporter for *hid* (*hid-GFP*), which contains the *hid* promoter but not coding region, together with GFP antibody labeling to assay p53 transcription factor activity (Figure 4A) (Tanaka-Matakatsu et al. 2009). Although GSCs did not apoptose in *p53^+^* (A+B+), they had high levels of hid-GFP expression after IR (Figure 4C) (Wylie et al. 2014; Xing et al. 2015; Ma et al. 2016). In contrast, IR did not induce high levels of hid-GFP in meiotic cells in region 2, suggesting that in these cells apoptosis is repressed upstream of proapoptotic gene expression (Figure 4C). Similar to the results for TUNEL, expression of the *hid-GFP* reporter in region 1 cystocytes was induced by IR in *p53^+^*(A+B+) wildtype and *p53^B41.5^* (A+B-) mutants, but not in *p53^5A-1-4^* (A-B-) null or *p53^A2.3^* (A-B+) mutants (Figure 4B-J). Together, these results indicate that, similar to the soma, p53A is necessary and sufficient for induction of proapoptotic gene expression and the apoptotic response to IR in the germline.

**Figure 4.**
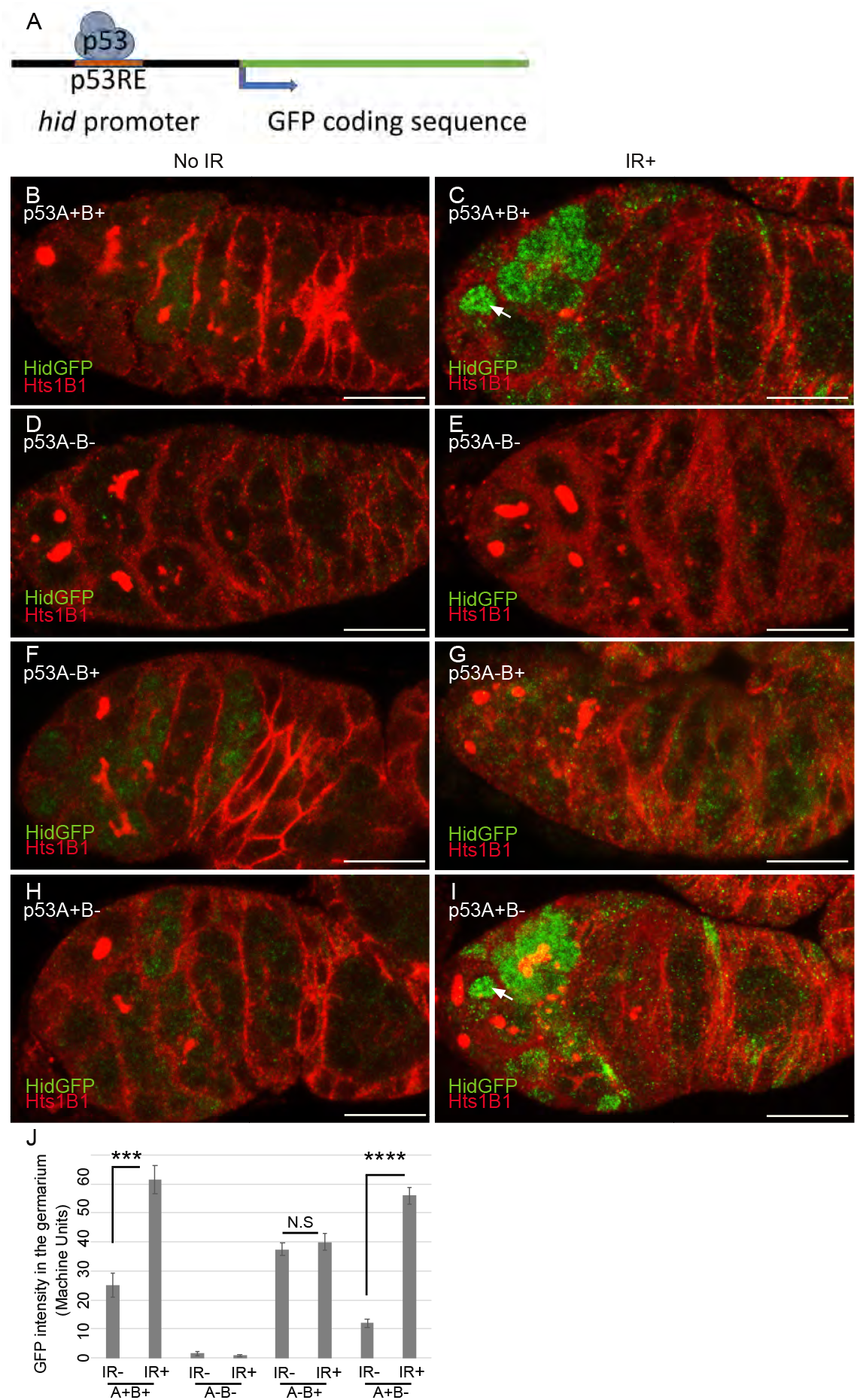
p53A is necessary and sufficient for IR-induced expression of proapoptotic genes in the germline. (**A**) Drawing of the *hid-GFP* promoter reporter which contains binding sites for p53 (Tanaka-Matakatsu et al. 2009). (**B-I**) *hid-GFP* expression without IR (**B, D, F, H**) or four hours after IR (**C, E, G, I**) in the germaria of *p53^+^* (A+B+) wild type (**B, C**), *p53^5A-1-4^* (A-B-) null (**D, E**), *p53^A2.3^* (A-B+) (**F, G**) and *p53^B41.5^* (A+B-) (**H, I**) females. Germaria were labeled for Hts (red) to identify GSCs and cytstocytes. Arrows in C, I indicate GFP positive GSCs. Scale bars are 10μm for **B-I**. (**J**) Quantification of the average GFP intensity in region 1 and 2 of the germaria for the genotypes shown in **B-I**. Averages are based on at least 20 germaria per genotype and two biological replicates. Error bars are S.E.M. ****: p<0.0001, ***: p<0.001 by unpaired Student’s t test.

### Both p53A and p53B isoforms respond to meiotic DNA breaks

The results indicated that although both p53A and p53B protein isoforms are expressed in the early germline, it is p53A that mediates the apoptotic response to IR in these cells. This result left unanswered what function p53B protein may have in the germline. A clue came when we noticed that there was a low level of *hid-GFP* expression beginning in late region 1 / early region 2a of the germarium even in the absence of IR (Figure 4B,D,F,H,J). This result is similar to one from the Abrams lab who found that in the absence of exogenous stress germline cells have low levels of expression of a promoter-reporter for another p53 target gene at the H99 locus, *reaper* (*rpr-GFP*), in germarium region 2, a time in oogenesis when meiotic DNA breaks are induced by the fly orthologue of Spo11 (Mei-W68) (Mehrotra and McKim 2006; Lu et al. 2010). They showed that this meiotic *rpr-GFP* expression was undetectable in either *mei-W68 or p53* null mutants, indicating that it was dependent on both meiotic DNA breaks and p53 (Lu et al. 2010).

To determine whether the p53A or p53B isoform responds to meiotic DNA breaks, we compared *hid-GFP* expression among the different *p53* alleles. To do this, we increased the exposure times for the unirradiated germaria from Figure 4, and replotted the quantification of their fluorescence intensity (Figure 5A-E). This clearly showed that in *p53^+^*(A+B+) wildtype germaria there is a low level of *hid-GFP* expression in all cells of a germline cyst in the absence of IR beginning in late region 1 / early region 2a, which was undetectable in the *p53^5A-1-4^* (A-B-) null mutant (Figure 5A,B, E). Thus, this *hid-GFP* expression in the absence of IR is dependent on p53 activity, similar to *rpr-GFP* that reports p53 activity in response to meiotic DNA breaks, (Lu 2010). In the *p53^B41.5^* (A+B-) mutant, *hid-GFP* expression was decreased to about 50% of wild type levels (Figure 5D,E). Given that *hid-GFP* expression was undetectable in the *p53* null, this result suggests that both p53A and p53B activate *hid-GFP.* Surprisingly, the *p53^A2.3^*(A-B+) mutant had higher levels of *hid-GFP* expression than wild type, and expression occurred earlier in oogenesis in region 1, including GSCs (Figure 5C, E). This result suggests that in the absence of the p53A isoform, the p53B isoform hyperactivates precocious expression of the *hid-GFP* reporter. This earlier *hid-GFP* expression in the *p53^A2.3^* (A-B+) mutant is not a response to meiotic DNA breaks, which are not induced until region 2a (Mehrotra and McKim 2006). Similar results were obtained when the isoform specific mutants were *in trans* to the *p53^5A-1-4^* null allele, indicating the results are not due to cryptic second-site mutations (data not shown). Altogether, these results suggest that both p53A and p53B are required for wild type levels of *hid-GFP* expression in the absence of IR. Given that p53 reporter expression in these cells is dependent on Mei-W68 (Spo11), these results further suggest that both isoforms respond to meiotic DNA breaks.

**Figure 5.**
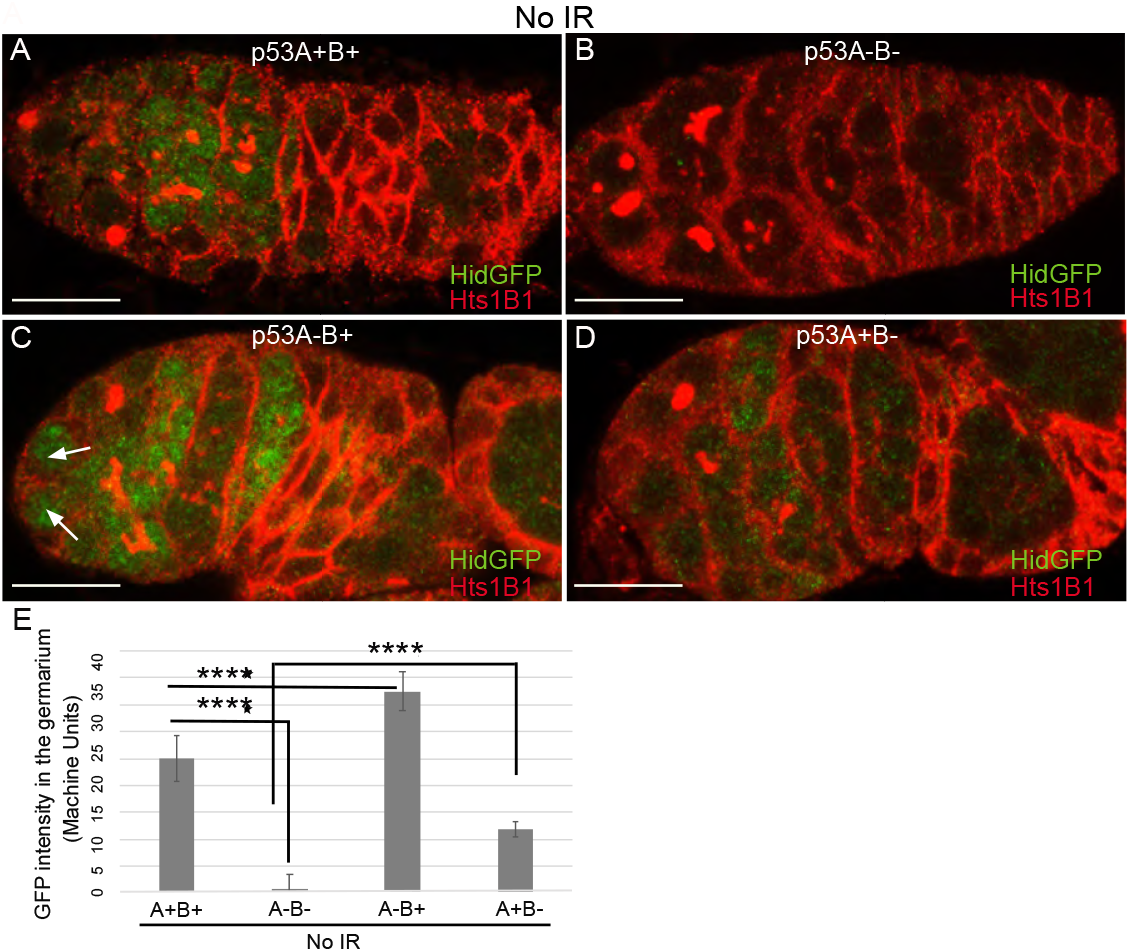
p53A and p53B both respond to meiotic DNA breaks. (**A-D**) Longer exposure micrographs of germaria from Figure 4 to show low level *hid-GFP* expression (green) without IR. Germaria were labeled with antibody against GFP (green) and Hts (red). Germarium from *p53^+^* (A+B+) wild type female (**A**) with *hid-GFP* expression that begins at the onset of meiosis in regions 2a-2b, which is undetectable in germaria from *p53^5A-1-4^* (A-B-) null females (**B**). The *p53^A2.3^* (A-B+) isoform mutant (**C**) had elevated and earlier expression of *hid-GFP*, including in GSCs (white arrows), whereas germaria from *p53^B41.5^* (A+B-) females (**D**) had *hid-GFP* expression that was 50% that of wild type. Scale bars in **A-D** are 10μm. (**E**) Quantification of the average GFP intensity in the germarium for the genotypes shown in **A-D**, replotted from Figure 4 J to compare *hid-GFP* intensity without IR among genotypes. Averages are based on 25 germaria per genotype and five biological replicates. Error bars are S.E.M. ****: p<0.0001.

### Dynamic p53B isoform abundance in p53 bodies correlates with meiotic DNA breaks

To investigate the relationship of p53 isoforms to meiotic DNA breaks further, we examined p53 isoform localization in the early germline. The level of GFP-p53A in p53 bodies was comparable among germline cells in all regions of the germarium, including GSC, dividing cystocytes in region 1, and during early stages of meiosis in regions 2a-2b (Figure 6A, A’). Ch-p53B was also abundant in p53 bodies in GSCs and most region 1 cystocytes (Figure 6B, B’). In contrast, the levels of p53B in the p53 bodies decreased at the onset of meiosis in late region 1 / early region 2, remained low in regions 2a-2b, and then increased again in most cells in late region 2b / early region 3 (Figure 6B, B’). Quantification of GFP-p53A and Ch-p53B levels within the same p53 bodies of GFP-p53A / Ch-p53B females confirmed that although Ch-p53B and GFP-p53A intensity in the bodies is approximately equal in region 1, p53B levels decrease to ~41% that of p53A in regions 2a-2b, and then increase again to levels comparable to p53A in regions 2b-3 (Figure 6C-G, Figure S4). This transient reduction in p53B levels in regions 2a-3 represents only ~24 hours of oogenesis, suggesting that it may be caused by relocalization of p53B protein from the p53 body to the nucleoplasm and then back again, rather than degradation and resynthesis of p53B protein (King 1970; Morris and Spradling 2011). The low magnitude and high variance of the nucleoplasmic fluorescence, however, precluded a determination as to whether the decrease of mCh-p53B in bodies is associated with a commensurate increase in the nucleoplasm (Figure 6E, S4 C-C’”). The transient decrease of p53B in bodies coincided with the known timing of meiotic DNA break induction in region 2a followed by subsequent DNA break repair by region 3. Moreover, p53B is required during this period to induce wild type levels of *hid-GFP*. These results suggest, therefore, that there may be a functional relationship between dynamic p53B localization and its response to meiotic DNA breaks.

**Figure 6.**
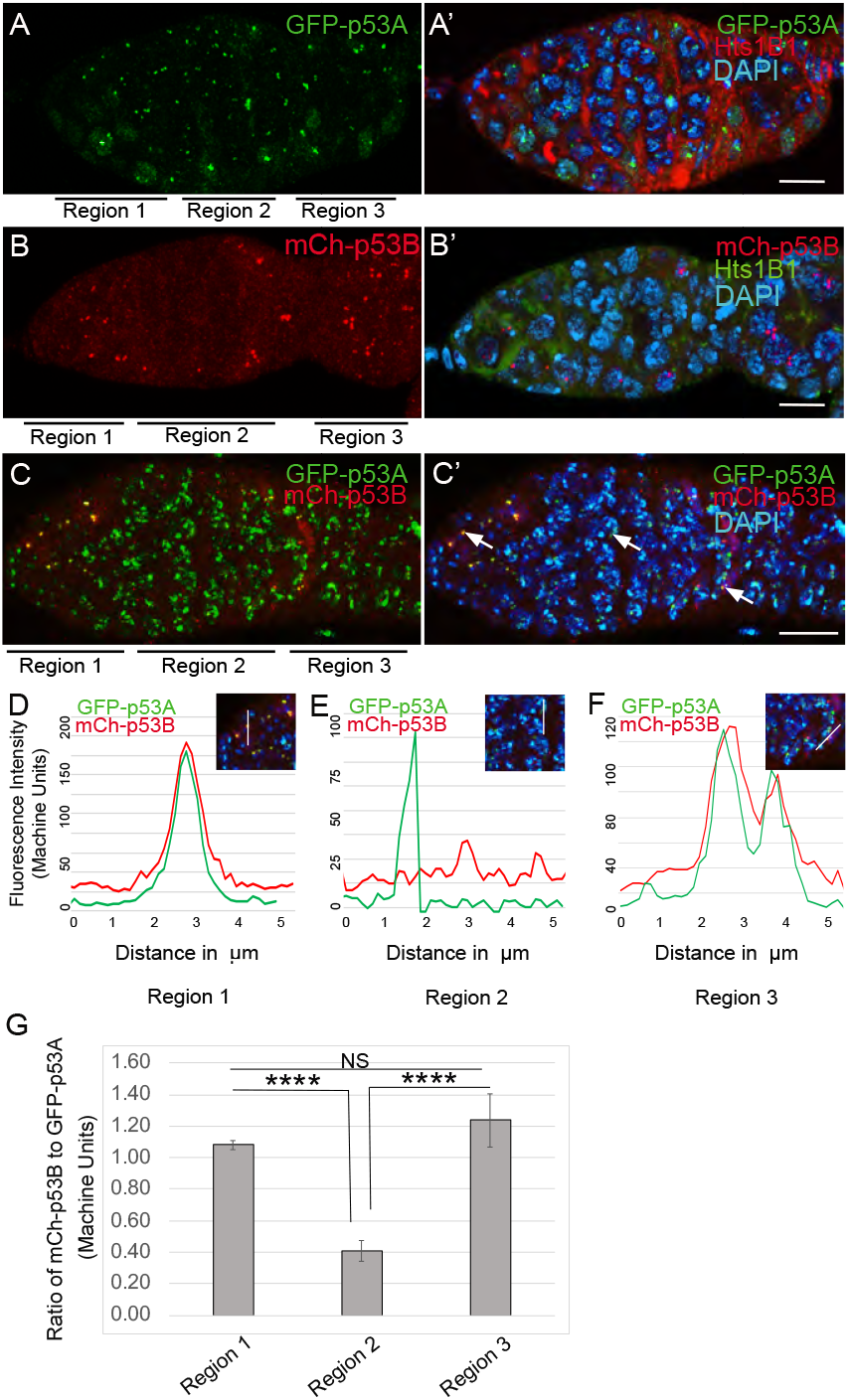
p53B protein levels fluctuate in p53 bodies during early meiosis. (**A-B’**) p53 bodies in germaria from GFP-p53A (**A, A’**, green) and mCh-p53B (**B, B’**, red) females. Germaria were labeled with antibodies against either GFP or mCherry, with Hts labeling in the other channel and DAPI labeling of nuclear DNA (blue) shown in **A’, B’**. Germarium regions are indicated below the images in **A-C**. Note that the intensity of mCh-p53B in the p53 bodies of germline nuclei was less at the onset of meiosis in region 2 than in regions 1 and 3. (**C, C’**). Colocalization of GFP-p53A and mCh-p53B in nuclear bodies in the germarium of a GFP-p53A / mCh-p53B female, with DAPI staining shown in C’. (**D-F**) Representative quantification of GFP-p53A and mCh-p53B fluorescent intensity within p53 bodies from region 1 (**D**), region 2 (**E**), and region 3 (**F**), indicated by white arrows in **C’**. Figures **A-B’** are single confocal z sections whereas **C-C’** are a composite stack of several z sections. Scale bars are 10μm. (**G**) Quantification of the ratio of mCh-p53B to GFP-p53A within bodies in gemarium regions 1, 2 and 3. Shown are mean and S.E.M.. **** = p< 0.0001 and n.s. = not significant by unpaired Student’s t test. n = 10 foci for regions 1 and 3 and 23 foci for region 2. See Figure S4 for more examples of quantification of p53A and p53B in nuclear bodies in regions 1-3.

### p53A and p53B mutants have persistent germline DNA damage

The *hid-GFP* results suggested that both p53A and p53B isoforms respond to meiotic DNA breaks. To investigate whether p53 isoforms regulate germline DNA breaks, we labeled ovaries with antibodies against phosphorylated histone variant H2AV (g-H2AV), which marks sites of DNA damage and repair, evident as distinct nuclear DNA repair foci (Madigan et al. 2002; Lake et al. 2013). It has been shown that labeling for g-H2AV detects repair foci at meiotic DNA breaks beginning in region 2a (Jang et al. 2003; Mehrotra and McKim 2006; Lake et al. 2013). Mei-W68 induces breaks in most cells of the 16-cell cyst, but g-H2AV repair foci are most abundant in four cells, one of which will become the oocyte while the others are destined to become nurse cells (Carpenter 1975; Jang et al. 2003; Mehrotra and McKim 2006; Lake et al. 2013; Hughes et al. 2018). Consistent with these previous reports, we observed that ovaries from wild type females had four cells per cyst with intense g-H2AV labeling, first appearing at the onset of meiotic recombination in germarium region 2a, decreasing in region 2b, and then undetectable in 97% of oocytes by germarium region 3 (oogenesis stage 1), a time when most meiotic DNA breaks have been repaired (Figure 7A, A’).

**Figure 7.**
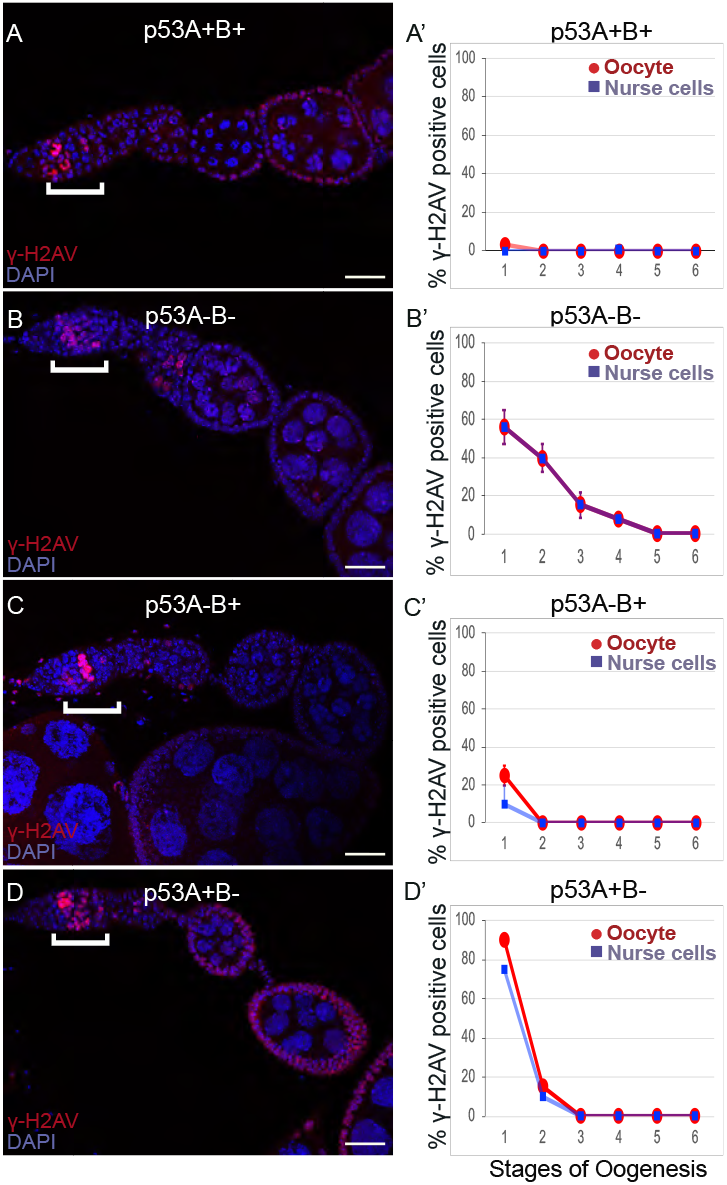
p53A and p53B mutants have persistent germline DNA damage. (**A-D**) Images of ovarioles of indicated genotypes immunolabeled with anti-g-H2Av (red) to detect DNA breaks, and counterstained with the DNA dye DAPI (blue), with corresponding quantification in **A’-D’**. The ovarioles are oriented with the anterior germarium to left (square bracket). Scale bars are 25μm. (**A’-D’**) Quantification of percent of ovarioles with g-H2Av positive nurse cells (blue squares) and oocyte (red ovals) at different stages in wild type *p53^+^* (A+B+) (**A’**), *p53^5A-1-4^* (A-B-) (**B’**), *p53^A2.3^* (A-B+) (**C’**), and *p53^B41.5^* (A+B-) (**D’**). Values are means of 2-3 biological replicates and > 20 ovarioles, with error bars representing S.E.M. Values with low variance have very small error bars that are not visible in the graphs. See supplemental Table I for *p* values.

In contrast, females homozygous mutant for the *p53^5A-1-4^* (A-B-) null allele had more than four germline cells per cyst with strong g-H2AV labeling (Figure 7B). This phenotype is similar to that of DNA repair mutants that increase the steady state number of unrepaired g-H2Av-positive breaks, and thereby the number of cells per cyst that label strongly for g-H2AV (Figure 7B) (Mehrotra and McKim 2006; Wei et al. 2019). Also similar to known DNA repair mutants, repair foci in *p53^5A-1-4^* persisted into later stages, with 56% of ovarioles having g-H2AV labeling in both nurse cells and oocytes in stage 1, and 8% of ovarioles having g-H2AV labeling as late as stage 4 (Figure 7B, B’, Table S1). Like the *p53^5A-1-4^* (A-B-) null, females homozygous for the *p53^A2.3^* (A-B+) allele also had more than four cells per cyst that labeled intensely for g-H2AV, as well as g-H2Av labeling in oocytes up to stage 1 in 25% of ovarioles, not as frequent as that in the *p53^5A-1-4^* (A-B-) null (56%) (Figure 7C, C’). The frequency of *p53^A2.3^* (A-B+) ovarioles with g-H2Av labeling in stage 1 was significantly different from wild type for the oocyte but not nurse cells (see Table S1 for p values). Females homozygous for *p53^B41.5^* (A+B-) also had more than four g-H2AV positive cyst cells and repair foci that persisted up to stage 2, later in oogenesis than in *p53^A2.3^* (A-B+) (stage 1), but not as late as in the *p53^5A-1-4^* (A-B-) null (stage 4) (Figure 7D, D’). g-H2AV foci were not observed in egg chambers after stage 6 in any genotype, suggesting that breaks are eventually repaired in the p53 mutants. The DNA damage in the *p53* mutant strains were at least as severe as those for mutants in genes required for meiotic recombination and repair, but we did not observe the egg ventralization or female sterility that have been described for those repair mutants (Ghabrial et al. 1998; Ghabrial and Schupbach 1999; Hughes et al. 2018). Altogether, these data reveal that p53A and p53B mutants have persistent DNA breaks in the *Drosophila* female germline.

### p53A and p53B isoforms are crucial for germline genome integrity when meiotic recombination repair is compromised

To further explore the role of *p53* isoforms in DNA break dynamics, we tested whether *p53* plays a more prominent role in the germline when there are defects in meiotic DNA recombination. We examined females doubly mutant for *p53* and *okra*, the fly ortholog of Rad54L, which is required for homologous recombination (HR) DNA repair in meiotic germline and somatic cells (Ghabrial et al. 1998; Sekelsky 2017; Hughes et al. 2018). It was previously reported that females doubly mutant for *okra* and a *p53* null allele result in egg chambers with extra nurse cells and shorter eggs, which was partially suppressed by mutation of *mei-W68*, but the relationship of this genetic interaction to DNA break repair, and the possible role of the different p53 isoforms, have not been explored (Lu et al. 2010).

Compared to wild type, females that were transheterozygous for two mutant *okra* alleles (*okra^RU/AA^*) had more cells per cyst with strong g-H2AV labeling in the germarium, which abnormally persisted into germarium region 3 (stage 1), consistent with previous reports that meiotic HR repair is delayed in *okra* mutants (Figure 8A-B’) (Jang et al. 2003; Mehrotra and McKim 2006; Lake et al. 2013). Also consistent with previous reports, *okra^RU/AA^* females were sterile (Ghabrial et al. 1998). Analysis of the different *okra;p53* double mutants indicated that they all had DNA break phenotypes that were more severe than the corresponding *p53* or *okra* single mutants. In *okra^RU/AA^*; *p53^5A-1-4^* (A-B-) double mutants, g-H2AV foci persisted up to stage 2 in 100% of ovarioles, with some ovarioles having repair foci in nurse cells and oocyte up to stage 5-6, ~30 hour later in oogenesis than wild type (Figure 8C, C’) (Lin and Spradling 1993). The *okra^RU/AA^* ; *p53^A2.3^* (A-B+) double mutants also had repair foci up to stage 6 in almost all ovarioles (Figure 8D, D’). Unlike *okra^RU/AA^*; *p53^5A-1-4^* (A-B-) null ovaries, however, repair foci in these *okra^RU/AA^*; *p53^A2.3^* (A-B+) were evident in the oocyte only (8D, D’). Given that *p53^A2.3^* expresses p53B but not p53A, these results suggest that in *okra* mutants the p53A isoform is required to protect genome integrity in the oocyte, while the p53B isoform is sufficient to rescue the DNA breaks in nurse cells. *okra^RU/AA^*; *p53^B41.5^* (A+B-) also had repair foci up to stage 6, but in these ovaries lacking p53B there were numerous repair foci in both the nurse cells and oocyte (Figure 8E, E’). Thus, p53B is required in nurse cells and oocytes. These data suggest that p53 isoforms are more crucial when HR is compromised, and that they have redundant and distinct functions in nurse cells and oocytes to protect genome integrity.

**Figure 8.**
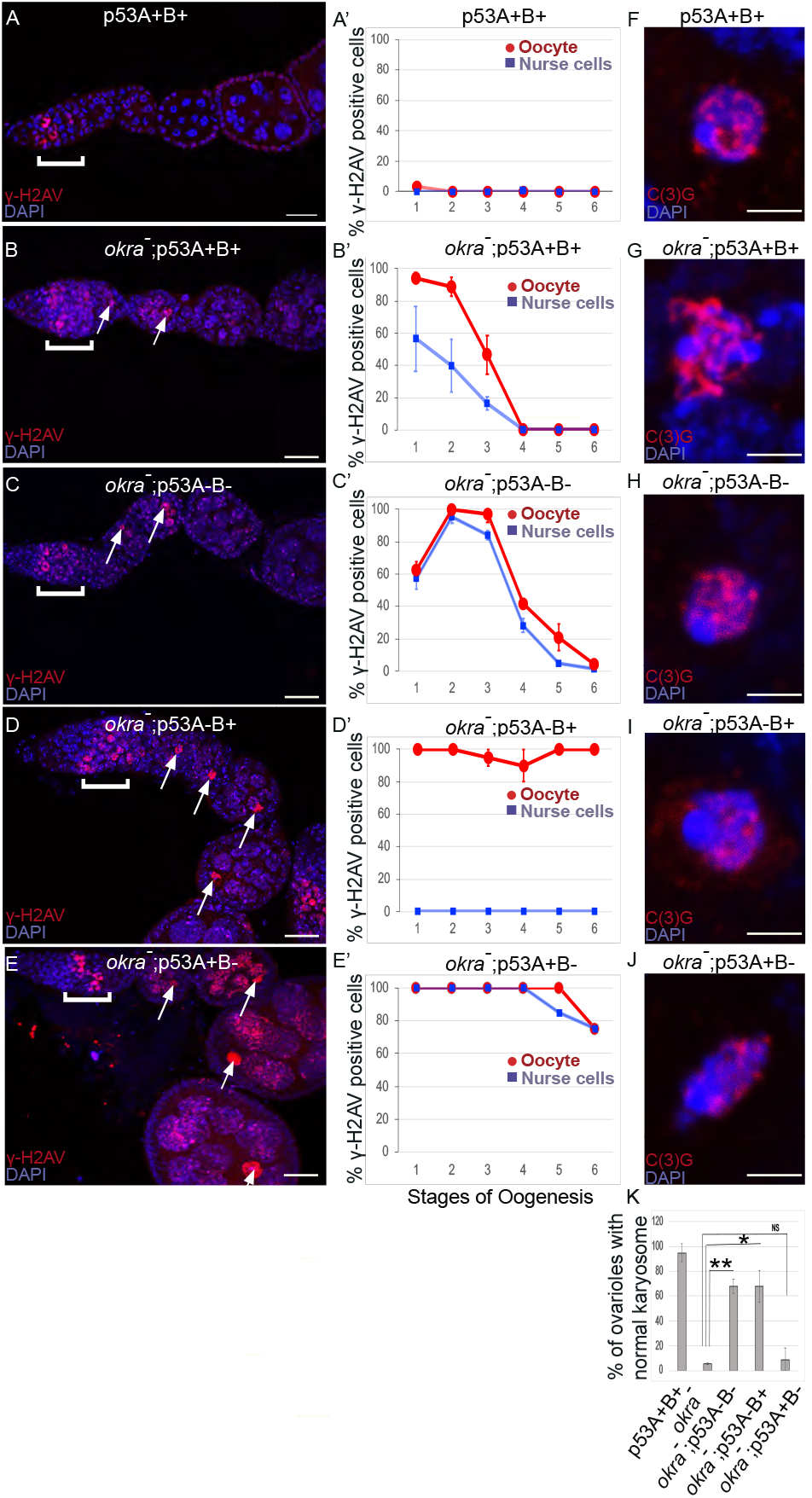
p53A and p53B have overlapping and distinct functions in germline genome integrity and the meiotic pachytene checkpoint. (**A-E**) *Drosophila* ovarioles of indicated genotypes were immunolabeled for g-H2Av (red) to detect DNA breaks and counterstained with the DNA dye DAPI (blue). The ovarioles are shown with the anterior germarium to left (square bracket). Scale bars are 25μm. (**A’-E’**) Quantification of percent of ovarioles with g-H2Av positive nurse cells (blue squares) and oocyte (red ovals) at different stages. (**A, A’**) *okra^+^; p53^+^* (A+B+) wild type. (**B, B’**) *okra^RU^ /okra^AA^* in a *p53^+^* (A+B+) wild type background. (**C, C’**) *okra^RU^ / okra^AA^; p53^5A-1-4^* (A-B-) null with delay in nurse cells and oocyte. (**D, D’**) *okra^RU^ / okra^AA^; p53^A2.3^* (A-B+) p53A mutant with repair foci in oocyte only. (**E, E’**) *okra^RU^ /okra^AA^; p53^B41.5^* (A+B-) p53B mutant with repair foci in both nurse cells and oocyte. Values in **A’-E’** are means of 2-3 biological replicates and > 20 ovarioles with error bars representing S.E.M. Those values that had low variance have very small error bars that are not visible in the graphs. See supplemental Table 1 for *p* values. (**F-K**) p53A is required for activation of the pachytene checkpoint. (**F-K**) Oocyte nuclei from stage 3-4 egg chambers labeled with antibodies against synaptonemal protein C(3)G (red) and DNA dye DAPI (blue). (**F**) Wild type with spherical compact karyosome. (**G**) *okra^RU^ / okra^AA^* with diffuse chromatin indicating activation of the pachytene checkpoint. (**H**) *okra^RU^ / okra^AA^; p53^5A-1-4^* (A-B-) null with compact spherical karyosome. (**I**) *okra^RU^ /okra^AA^; p53^A2.3^* (A-B+) p53A mutant with spherical karyosome. (**J**) *okra^RU^ / okra^AA^; p53^B41.5^* (A+B-) with elipitical nucleus. Scale bars are 3μm. (**K**) Quantification of karyosome formation. Data are means based on two biological replicates with ~30 nuclei per strain per replicate, with error bars representing S.E.M. * p < 0.05, ** p < 0.01, n.s. = not significant by unpaired Student’s t test.

### p53A is required for the meiotic pachytene checkpoint

Defects in DNA recombination and persistent DNA damage are known to activate a meiotic pachytene checkpoint arrest in multiple organisms (Bahler et al. 1994; Gebel et al. 2017). During oogenesis in mice and humans, the pachytene arrest is mediated by p63 and p53 (Suh et al. 2006; Bolcun-Filas et al. 2014; Coutandin et al. 2016; Gebel et al. 2017; Marcet-Ortega et al. 2017; Rinaldi et al. 2020). In *Drosophila*, it is known that defects in the repair of meiotic DNA breaks activate the pachytene checkpoint, but it is not known whether this checkpoint response requires p53 (Ghabrial et al. 1998; Joyce and McKim 2011; Hughes et al. 2018). To address this question, we took advantage of a visible manifestation of the *Drosophila* pachytene checkpoint arrest, which is a failure of the oocyte nucleus to form a compact spherical structure known as the karyosome, which normally occurs during stage 3 of oogenesis (Ghabrial et al. 1998). We examined karyosome formation by labeling with antibodies against the synaptonemal complex (SC) protein C(3)G and DAPI (Page and Hawley 2001). In wild type females, 95% of ovarioles had a spherical compact karyosome beginning in stage 3 (Figure F, K). In *okra^RU/AA^* mutants, however, compaction was normal in only 5% of ovarioles, with the oocytes in 95% of ovarioles appearing either diffuse or ellipsoidal, morphologies that have been described in previous studies that showed that the DNA repair defects in *okra* mutants strongly activate the pachytene checkpoint (Figure 8 G, K) (Ghabrial et al. 1998). In *okra^RU/AA^; p53^5A-1-4^* (A-B-) null double mutants, however, karyosome compaction was normal in 68% of ovarioles, a fraction that is significantly different than the *okra* single mutant, suggesting that *p53* is required for normal activation of the pachytene checkpoint (Figure 8H, K). Similarly, in *okra^RU/AA^; p53^A2.3^* (A-B+) double mutants karyosome compaction was normal in 68% of ovarioles, significantly different than the fraction in *okra* single mutants alone (Figure 8I, K). This failure to activate the pachytene checkpoint in the majority of *okra^RU/AA^;p53^A2.3^* (A-B+) ovarioles is even more striking given that they had a more severe DNA break phenotype in the oocyte than did *okra* single mutants that strongly engaged the checkpoint (Figure 8B, B’, D, D’). In contrast, *okra^RU/AA^; p53^B41.5^* (A+, B-) mutants had normal karyosome compaction in only 9% of ovarioles, a fraction that was not significantly different than *okra* single mutants, indicating that the pachytene checkpoint was activated in most of these ovarioles that expressed p53A but not p53B (Figure 8J, K). Altogether these data suggest that the p53A isoform is required for normal pachytene checkpoint activation when meiotic DNA recombination is impaired, analogous to the functions of mammalian p53 and p63.

## Discussion

A common property of the p53 gene family across organisms is that they encode multiple protein isoforms, but their functions are largely undefined. We found that the *Drosophila* p53B protein isoform is more highly expressed in the germline where it colocalizes with a shorter p53A isoform in subnuclear bodies. Despite this p53B germline expression, it is the p53A isoform that was necessary and sufficient for the apoptotic response to IR in both the germline and soma. In contrast, p53A and p53B both responded to programmed meiotic DNA breaks. Although apoptosis is repressed in meiotic cells, p53A and p53B mutants had persistent DNA breaks, a phenotype that was enhanced when meiotic DNA recombination was defective. The role of the p53 isoforms to prevent persistent DNA breaks was cell type specific, with p53B being necessary and sufficient in nurse cells, whereas both p53B and p53A were required in the oocyte. Our data has also uncovered a requirement for the *Drosophila* p53A isoform in the meiotic pachytene checkpoint response to unrepaired DNA breaks. Overall, these data suggest that *Drosophila* p53 isoforms have evolved overlapping and distinct functions to mediate different responses to different types of DNA damage in different cell types. These findings are relevant to understanding the evolution of p53 isoforms, and have revealed interesting parallels to the function of mammalian p53 family members in oocyte quality control.

### p53 localizes to subnuclear bodies in *Drosophila* and humans

p53 isoforms colocalized to subnuclear bodies in the *Drosophila* male and female germline (Figure 9A). This finding is consistent with a previous study that reported p53 bodies in the *Drosophila* male germline, although that study did not examine individual isoforms (Monk et al. 2012). We deem it likely that these p53 bodies form by phase separation, an hypothesis that remains to be formally tested (Mitrea and Kriwacki 2016; Alberti 2017). *Drosophila* p53 subnuclear bodies are reminiscent of human p53 protein localization to subnuclear PML bodies (Mauri et al. 2008). Evidence suggests that trafficking of p53 protein through PML bodies mediates p53 post-translational modification and function, although the relationship between nuclear trafficking and the functions of different p53 isoforms has not been fully evaluated (Fogal et al. 2000; Chang et al. 2018). Similarly, we observed a decline in abundance of p53B within p53 bodies in germarium region 2a, followed by a restoration of p53B within bodies in region 3. This fluctuation of p53B in bodies temporally correlates with the onset of meiotic DNA breaks in region 2a and their repair in regions 2b - 3. Although p53B was undetectable in p53 bodies in many cells of region 2, the analysis of the p53B mutant alleles indicated that it is required in those cells to activate a *hid-GFP* reporter to wild type levels and to prevent persistent DNA breaks. These observations are consistent with the idea that nuclear trafficking of p53B out of bodies may mediate its response to meiotic breaks, although further experiments are required to test this idea (Figure 9A). Future analysis of *Drosophila* p53 bodies will help to define how p53 isoform trafficking mediates the response to genotoxic and other stresses.

**Figure 9.**
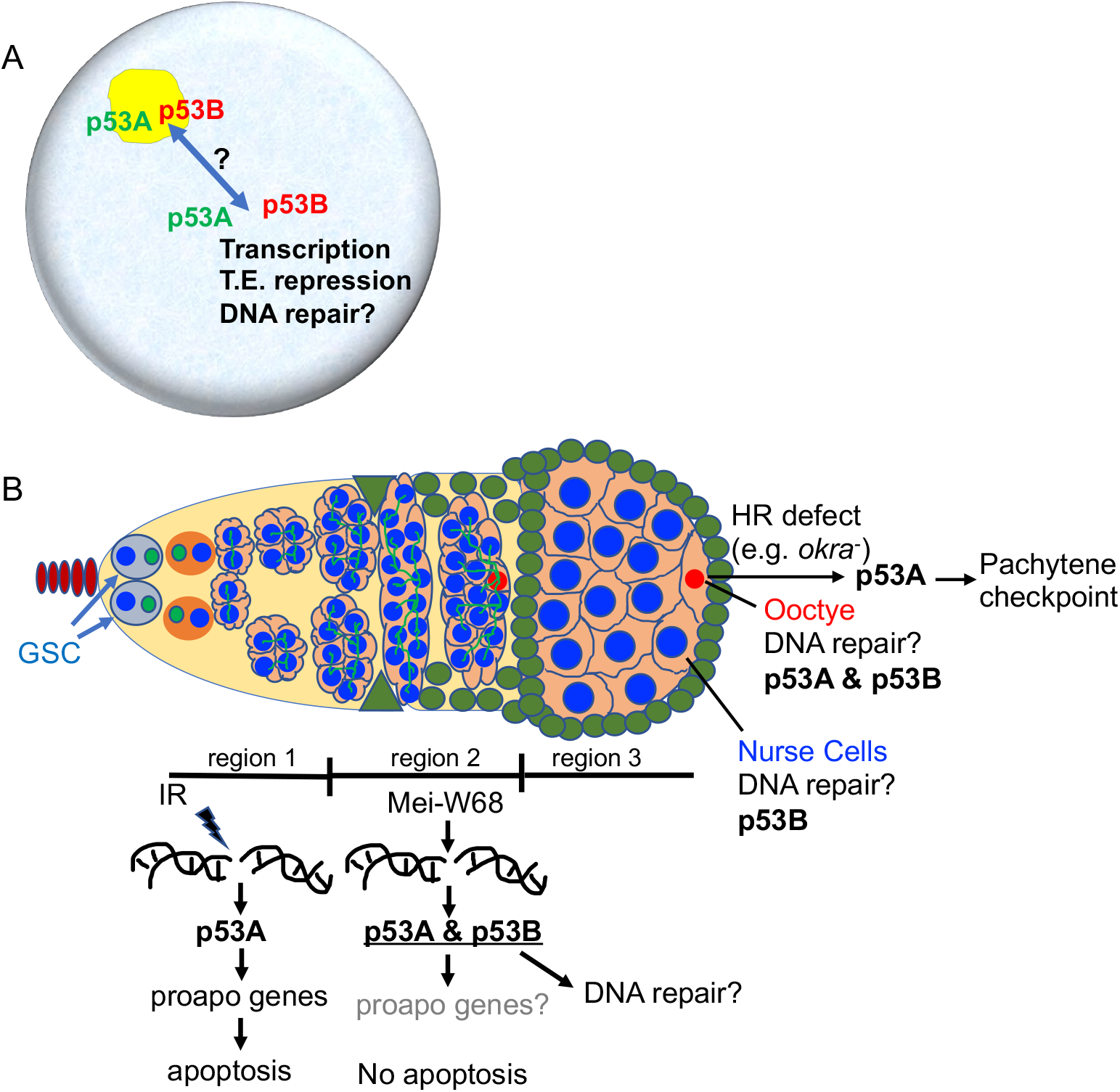
Model: Drosophila p53 isoforms colocalize to nuclear bodies and have DNA lesion and cell type specificity in the germline genotoxic stress response. **(A)** The p53A (green) and p53B (red) isoforms are concentrated in p53 bodies of germline nuclei (blue). Trafficking of p53 isoforms in and out of these bodies (double arrow) may mediate their functions in transcription, transposable element (T.E.) repression, and DNA repair. (**B**) The p53A isoform mediates the apoptotic response to IR in dividing germline cells in region 1 of the germarium. This apoptotic response is repressed in germline stem cells (GSCs) and meiotic cells. Both the p53A and p53B isoforms are activated by mei-W68-induced meiotic breaks. The data lead to the proposal that p53B is required for repair of meiotic DNA breaks in nurse cells (blue) and oocytes (red) nuclei, whereas p53A is required for DNA repair and activation of the meiotic pachytene checkpoint in oocytes when homologous recombination (HR) is defective.

### p53A mediates the apoptotic response to IR in the soma and germline, whereas both p53A and p53B respond to meiotic DNA breaks

Our data indicate that the p53A isoform is necessary and sufficient for activation of proapoptotic gene expression and apoptosis in response to IR, in both the soma and germline. Alleles that expressed only p53A had IR-induced *hid-GFP* expression and apoptosis that was equivalent to wild type, whereas those that expressed only p53B were equivalent to p53 null alleles. While our manuscript was in preparation, it was reported that p53A and p53B both participate in the apoptotic response to IR in the ovary (Park et al. 2019). That study used the GAL4 / UAS system to express either p53A or p53B rescue transgenes in a *p53* null background. In contrast, we created and analyzed loss-of-function, isoform-specific alleles at the endogenous *p53* locus, which we believe more accurately reflect the physiological function of p53 isoforms. We favor the conclusion, therefore, that it is the p53A isoform that has the exclusive function of mediating the apoptotic response to IR in the soma and germline.

Dividing germline cystocytes in germarium region 1 apoptosed after IR, but their ancestor GSCs and descendent meiotic cells did not (Figure 9B). The expression of the *hid-GFP* promoter reporter in GSCs is consistent with previous evidence that apoptosis is repressed in these stem cells downstream of *hid* transcription by the miRNA *bantam* (Wylie et al. 2014; Xing et al. 2015; Ma et al. 2016). How meiotic cells repress apoptosis is not known, although it is crucial that they do so because they have programmed DNA breaks. Our finding that IR does not induce high levels of *hid-GFP* in meiotic cells suggests that, unlike GSCs, apoptosis is repressed in meiotic cells upstream of p53-regulated proapoptotic gene expression.

Although p53B did not mediate the apoptotic response to IR, our data indicate that both p53B and p53A respond to meiotic DNA breaks (Figure 9B). In the absence of IR, we observed p53-dependent expression of the *hid-GFP* reporter during early meiosis, consistent with previous evidence that p53 is activated by meiotic DNA breaks (Lu et al. 2010). While *hid-GFP* expression was undetectable in *p53* null mutants, the p53B mutant alleles, which express p53A, reduced expression by only 50%. Together, these results suggest that both p53A and p53B respond to meiotic DNA breaks. The results for the p53A mutant allele were not informative for this hypothesis, however, because in that mutant *hid-GFP* expression was higher earlier in oogenesis. The mechanism for this gain of function phenotype is not clear, but a cogent hypothesis is that the p53A subunit restrains the transcription factor activity of the more potent p53B in heterotetramers, and in the p53A isoform mutants the exclusive formation of more active p53B homotetramers induces higher and precocious *hid-GFP* expression. This hypothesis is consistent with our previous evidence that the p53B isoform with a longer TAD is a much stronger transcription factor than p53A, and that p53A and p53B can form heterocomplexes (Zhang et al. 2015). Moreover, it is known that in humans shorter p53 isoforms repress the transcriptional activity of longer ones in heterotetramers (Anbarasan and Bourdon 2019). It is important to note, however, that while *hid* expression was constitutively higher in the p53A mutants than in wild type, it was lower than that induced by IR, and did not result in apoptosis. Altogether, the data indicate that p53A responds to DNA damage caused by IR, whereas both p53A and p53B respond to programmed meiotic DNA breaks. Important questions motivated by our results are how these distinct responses to DNA damage are determined by different types of DNA lesions, checkpoint signaling pathways, and p53 isoform structure.

### p53 isoforms have redundant and distinct requirements to prevent germline DNA damage

*p53* mutants had persistent DNA breaks in the *Drosophila* female germline (Figure 9B). These persistent germline DNA breaks may reflect a requirement for p53 for the timely repair of meiotic DNA breaks, and / or the prevention of new breaks by other mechanisms. Consistent with a role in meiotic DNA break repair, *p53* mutants had an increased number of cells with g-H2AV foci beginning in germarium stage 2a, a time when mei-W68 induces programmed meiotic double strand DNA breaks. Moreover, g-H2AV foci were continuously observed to different later times in oogenesis depending on the *p53* genotype, which was enhanced by *okra* (RAD54L) mutations that are known to compromise the repair of meiotic DNA breaks. These observations are consistent with the idea that meiotic DNA break repair is delayed for different amounts of time in the different *p53* genotypes. It has also been shown, however, that *p53* is required with the piRNA pathway to fully restrain mobile element activity in the female germline (Wylie et al. 2016; Wei et al. 2019). A non-mutually exclusive possibility, therefore, is that the increase in germline DNA breaks in the *p53* mutants is also the result of elevated mobile element activity.

The proposed role for *Drosophila* p53 in the repair of meiotic DNA breaks is consistent with evidence from other organisms that p53 has both global and local roles in DNA repair. It is known that *Drosophila* p53 and specific isoforms of human p53 induce the expression of genes required for different types of DNA repair (Brodsky et al. 2004; Gong et al. 2015; Williams and Schumacher 2016). p53 also acts locally at DNA breaks in a variety of organisms, including humans, where it can mediate the choice between HR versus non-homologous end joining (NHEJ) repair (Moureau et al. 2016; Williams and Schumacher 2016). In fact, it has been shown that human p53 directly associates with RAD54 at DNA breaks to regulate HR repair, consistent with our finding that *p53; okra* (RAD54L) double mutants have enhanced DNA repair defects (Linke et al. 2003). Moreover, the the *C. elegans* p53 ortholog CED-4 locally promotes HR and inhibits NHEJ repair in the germline (Mateo et al. 2016). Thus, we deem it likely that the DNA damage that we observe in the germline of *Drosophila p53* mutants may, in part, reflect a local requirement for p53 protein isoforms to regulate meiotic DNA repair (Figure 9A, B).

Our analysis also revealed that p53 isoforms have overlapping and distinct requirements to prevent persistent DNA damage in different cell types (Figure 9B). The results indicated that both p53A and p53B are required in the oocyte, whereas p53B is necessary and sufficient in nurse cells, even though nurse cells express both p53A and p53B isoforms. This differential requirement for p53 isoforms may reflect differences in how meiotic breaks are repaired in nurse cells versus oocytes. While it is not known whether DNA repair pathways differ between nurse cells and oocytes, evidence suggests that the creation of meiotic breaks does, with breaks in pro-oocytes but not pro-nurse cells depending on previous SC formation (Mehrotra and McKim 2006). Important remaining questions include whether different p53 isoforms participate in, and influence the choice among, different DNA repair pathways in different cell types.

### Similar to the mammalian p53 family, *Drosophila* p53A is required for the meiotic pachytene checkpoint

Our study has also uncovered a requirement for *Drosophila p53* in the meiotic pachytene checkpoint. This function was isoform specific, with p53A, but not p53B, being required for full checkpoint activation in oocytes with persistent DNA breaks. The failure to engage the pachytene checkpoint in the majority of *okra;p53^A2.3^* double mutant oocytes is more striking given that these cells had more severe DNA repair defects than the *okra* single mutants that strongly engaged the checkpoint. Some *okra; p53* null egg chambers had a pachytene arrest, which suggests that p53-independent mechanisms also activate the checkpoint, perhaps in response to secondary defects in chromosome structure which are known to independently trigger the pachytene checkpoint in flies and mammals (San-Segundo and Roeder 1999; Wu and Burgess 2006; Li and Schimenti 2007; Joyce and McKim 2009). Moreover, although the pachytene checkpoint was strongly compromised in the p53 null and p53A mutant alleles, they did not suppress *okra* female sterility, suggesting that other mechanisms ensure that eggs with excess DNA damage do not contribute to future generations. Altogether, the results indicate that p53A is required for both DNA repair and full pachytene checkpoint activation in the oocytes.

Evidence suggests that the ancient function of the p53 family was of a p63-like protein in the germline (Levine 2020). Consistent with this, our findings in *Drosophila* have parallels to mammals where the TAp63α isoform and p53 mediate a meiotic pachytene checkpoint arrest, and the apoptosis of millions of oocytes that have persistent defects (Di Giacomo et al. 2005; Suh et al. 2006; Bolcun-Filas et al. 2014; Gebel et al. 2017; Rinaldi et al. 2017; Rinaldi et al. 2020). Our evidence suggests that the different isoforms of the sole *p53* gene in *Drosophila* may subsume the functions of vertebrate p53 and p63 paralogs to protect genome integrity and mediate the pachytene arrest. Unlike p53 and p63 in mammals, however, *Drosophila* p53 does not trigger apoptosis of defective oocytes. Instead, the activation of the pachytene checkpoint disrupts egg patterning, resulting in inviable embryos that do not contribute to future generations (Hughes et al. 2018). Thus, in both *Drosophila* and mammals, the p53 gene family participates in an oocyte quality control system that ensures the integrity of the transmitted genome.

## Materials and Methods

### Drosophila genetics

Fly strains were reared at 25 C. Fly strains were obtained from the Bloomington Drosophila Stock Centre (BDSC, Bloomington, IN, USA) unless otherwise noted. The *hid-GFP* fly strain was a generous gift from W. Du (Tanaka-Matakatsu et al. 2009). A *y w^c67C23^* strain was used as wild type. The *okra* mutant strains were obtained from Trudi Shupbach’s lab. See Table S2 for a complete list of fly strains used in this study.

### Creation and characterization of p53A and p53B isoform specific alleles

Isoform specific alleles of *p53* were generated by injection of gRNA encoding plasmid into embryos of a nanos-Cas9 strain using standard methods (Gratz et al. 2013; Ren et al. 2013). Candidate lines for p53B-specific alleles were initially identified by screening for co-knockout of *white* (Ge et al. 2016). Injections were performed by Rainbow Transgenics (USA). Alleles were identified by PCR genotyping. The *p53^A2.3^* allele is a 23 bp deletion (coordinates 23053346–23053368 in the *D. melanogaster* genome version 6.32), and has a 7 base-pair (Robin et al. 2019). The *p53^B41.5^* allele is a 14 bp deletion (coordinates 23053726-23053739) with an insertion of a single Adenine. RT-PCR analysis of *p53* isoform-specific mutants was performed on mRNA from adult flies using standard methods. See Figure S3 and Table S2 for further information about alleles, gRNAs and primers.

### Gamma irradiation and cell death assays

Adult females were mated and conditioned on wet yeast for three days. They were then irradiated with a total of 4000 rad (40 Gy) from a Cesium source and were allowed to recover at 25 C for 4 hours before TUNEL labeling. TUNEL labeling (In Situ cell death detection kit, Fluorescein, Roche) was performed according to manufacturer’s instructions. Follicle cell death in Figure 2 was quantified by counting cells TUNEL positive cells in stage 6.

### Immunofluorescent microscopy

Dissection, fixation, antibody labeling and immunofluorescent microscopy of testes and ovaries were as previously described (Thomer et al. 2004). Primary antibodies and concentrations used were rabbit anti-GFP (Invitrogen) 1:500, rabbit anti-dsRed (Clontech) 1:200, mouse anti dsRed (Clontech) 1:200, mouse anti Hts1B1 (DSHB) 1:20 (Zaccai and Lipshitz 1996), and mouse anti-gH2AV (DSHB) 1:1000 (Lake et al. 2013). The anti-gH2AV antibody was preabsorbed against fixed wild type ovaries before use. Secondary antibodies were Alexa 488 anti-rabbit, Alexa 488 anti-mouse, Alexa 568 anti-rabbit, and Alexa 568 anti-mouse (Jackson) all used at 1:500-1-750. Samples were counterstained with DNA dye 4’,6-diamidino-2-phenylindole (DAPI) at 1μg / ml. Confocal micrographs were captured on a Leica SP8 confocal using a 63X multi-immersion lens.

### Fluorescent quantification

Hid-GFP fluorescence in Figures 3 and 4 was quantified using the LASX software of the Leica Sp8 confocal microscope. Total GFP intensity was measured across a composite z-stack of each germarium. See Figure legends for sample sizes, biological replicates, and p values.

The intensities of GFP-p53A and mCh-p53B in p53 bodies in Figures 1, 6, and S4, were quantified along a line using LASX software on the Leica SP8. For strains expressing both GFP-p53A and mCh-p53B, the ratio of GFP-p53A: mCh-p53B was quantified within each body. For those expressing a single tagged isoform, the ratios of intensities in germarium regions 1:2 or 2:3 among different bodies was calculated, all within the same germarium to control for technical variation. See figure legends for sample sizes and p values.

g-H2Av was quantified in Figures 7 and 8 by scoring cells whose total fluorescent intensity was above the threshold of wild type controls. Nurse cells and oocytes were scored as positive or negative in different stage egg chambers within each ovariole. See Figure legends and Table S1 for sample sizes and p values.

## Acknowledgements

We thank W. Du, K. McKim, T. Shupbach and the Bloomington Drosophila Stock Center for fly strains; S. Hawley and J. Sekelsky for antibodies; and FlyBase for critical information. Thank you to H. Herriage, S. Brown, and M. Dykhouse for helpful discussions. Thanks to J. Powers of the IU Light Microscopy Imaging Center (LMIC) for imaging advice and support. This research was supported by NIH 2R01GM113107 to B.R.C.

## Supplemental Data

**Figure S1.**
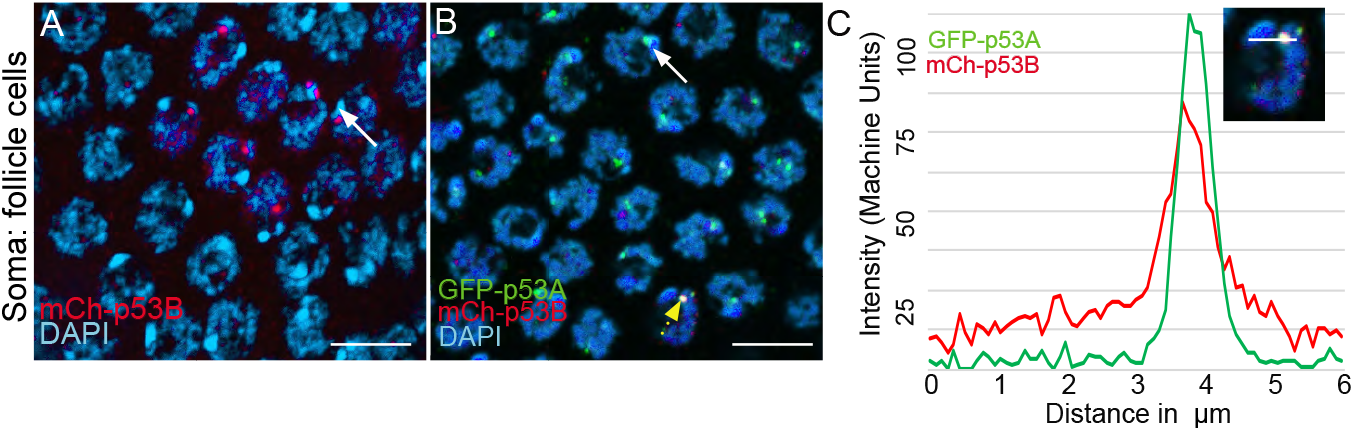
Rare Ch-p53B expression in somatic follicle cells. (**A, B**) mCh-p53B in nuclear bodies in a small group of follicle cells in a mCh-p53B female (**A**), and colocalization with GFP-p53A in follicle cells in a mCh-p53B / GFP-p53A female (**B**). White arrows indicate two examples of DAPI-bright pericentric heterochromatin, near which p53 bodies are often located. (**C**) Quantification of GFP-p53A and mCh-p53B labeling in one nuclear body in **B** (dotted yellow arrow), measured along a 6 μm line (inset). Scale bar is 10μm.

**Figure S2.**
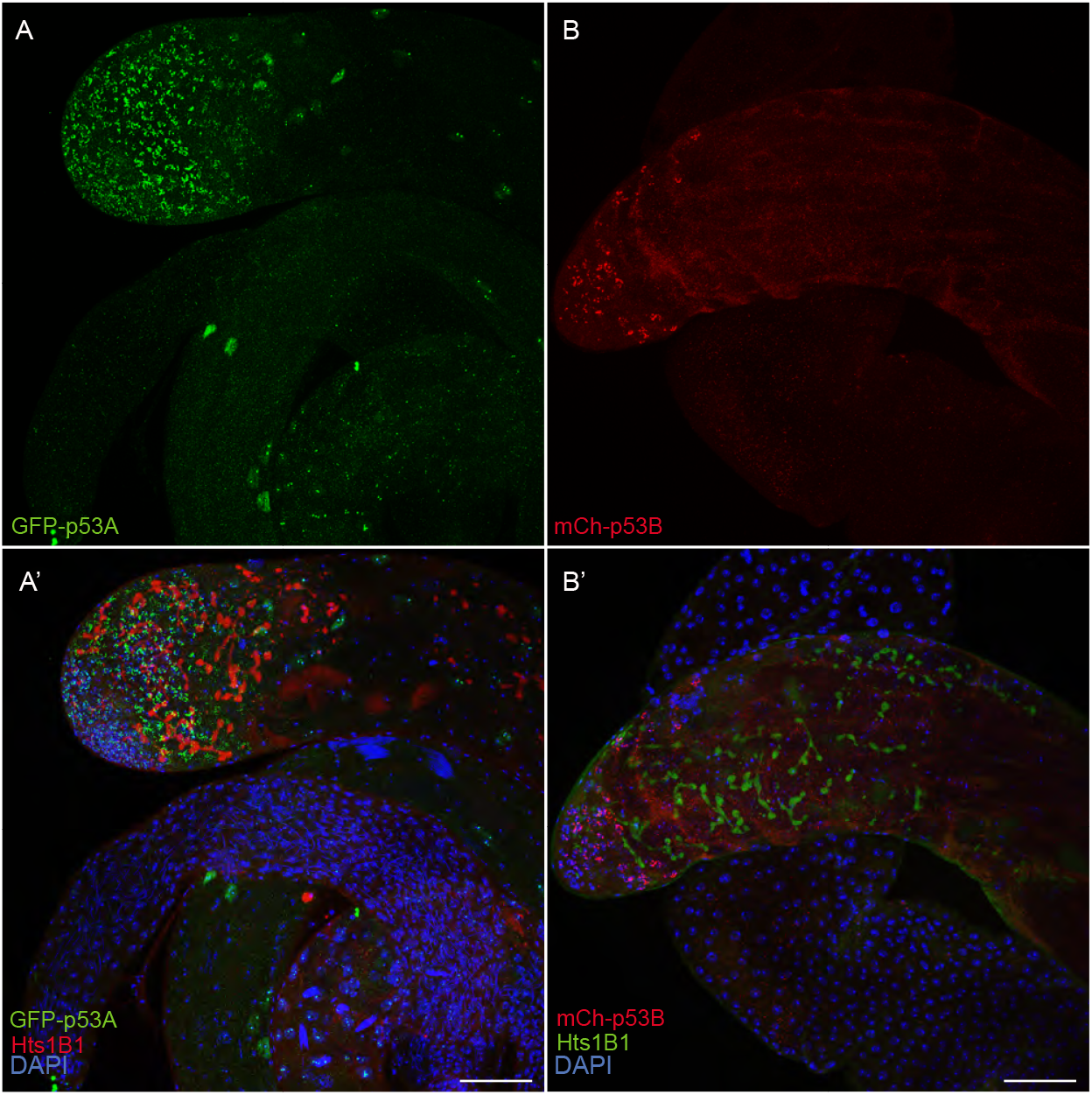
p53A and p53B are expressed in the male germline. A testis expressing GFP-p53A (**A, A’**) or mCh-p53B (**B, B’**), with labeling with anti-Hts (germline fusomes) and DAPI (blue) shown in **A’B’**. Scale bar is 20μm

**Figure S3.**
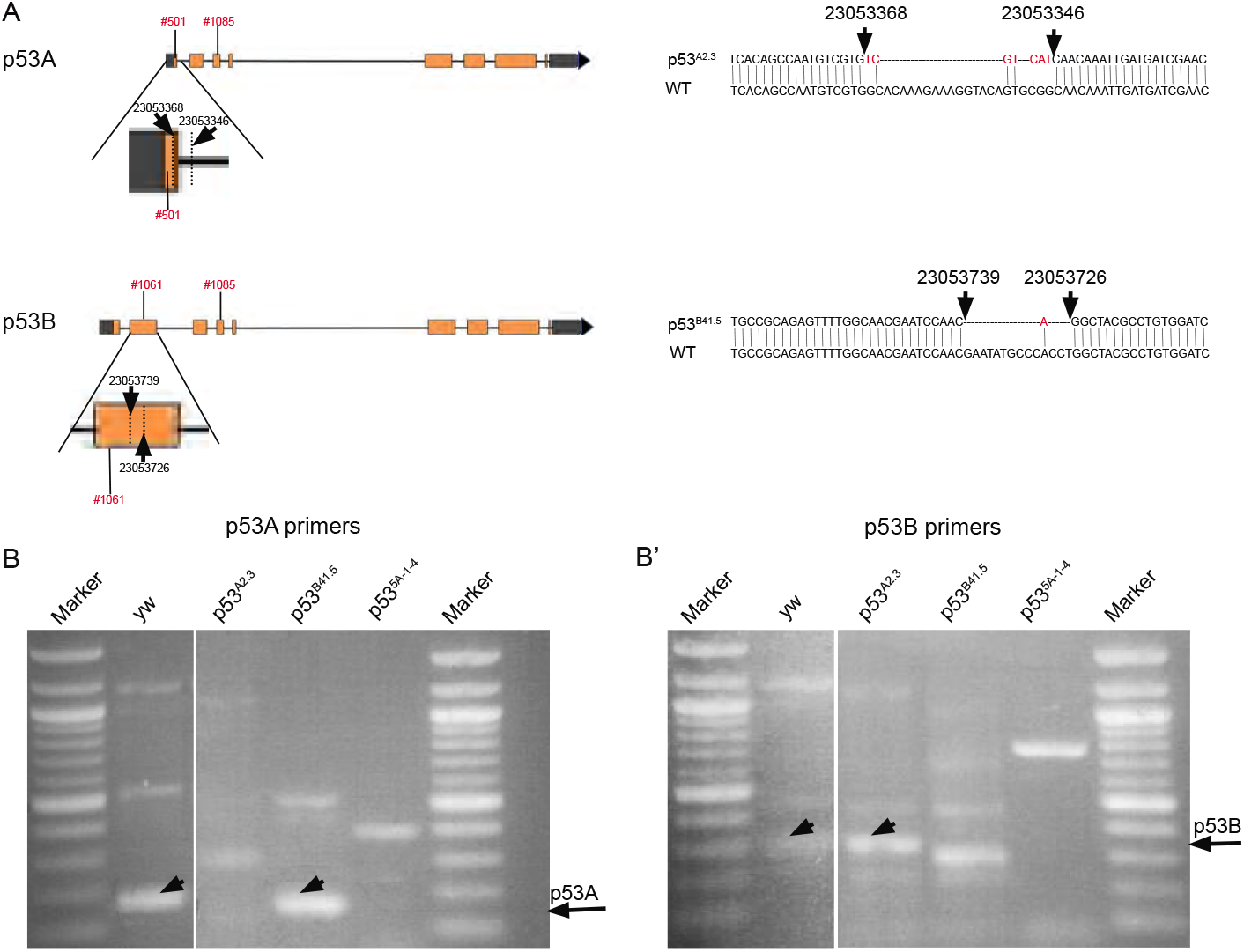
*p53* isoform-specific alleles. (**A**) Left: Shown are the location of *p53^A2.3^* and *p53^B41.5^* deletion alleles and the coordinates of primers (red font) used for RT-PCR. Expanded maps below indicate the coordinates of the deletion endpoints within the p53A and p53B mRNA isoforms (dotted lines and straight arrows). Right: DNA Sequence of *p53^A2.3^* and *p53^B41.5^* deletion alleles (top) aligned with *Drosophila* reference genome sequence (WT, bottom). Red nucleotides indicate unique nucleotide insertions. All genomic coordinates are based on *Drosophila melanogaster* genome release 6.32. (**B, B’**) RT-PCR with the primer pairs indicated in **A** that are specific to p53A mRNA (**B**) or p53B mRNA (**B’**) from *p53^+^* (A+B+), *p53^A2.3^* (A-B+), *p53^B41.5^* (A+B-), *p53^5A-1-4^* (A-B-). Black arrows indicate position of p53A and p53B PCR products.

**Figure S4.**
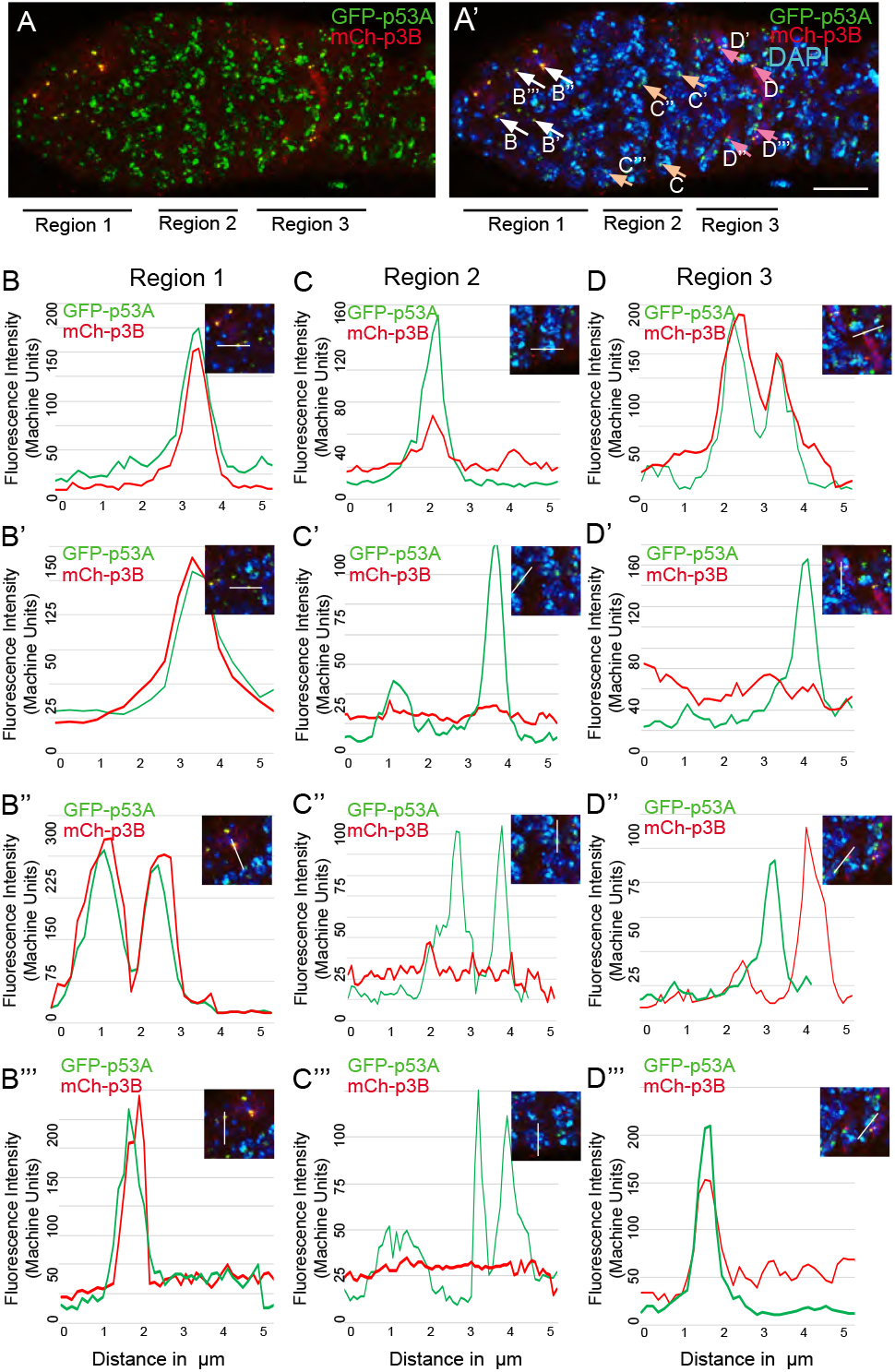
mCh-p53 levels fluctuate in p53 bodies during early meiosis. (**A, A’**) A germarium from a GFP-p53A / mCh-p53B female showing localization of p53A and p53B proteins to p53 nuclear bodies in germarium regions 1, 2 and 3, with DAPI labeling of DNA (blue) shown in **A’**. Arrows in **A’** indicate the bodies within which GFP-p53A and mCh-p53B were quantified and the letters within the figure indicate the corresponding panels shown below. Scale bar is 10μm. (**B-D**”‘) Four examples each of fluorescent quantification along a 6 μm line within nuclear bodies from regions 1 (**B-B”‘**), 2 (**C-C”‘**), and 3 (**D-D”‘**), which are indicated by arrows in **A’**. Related to figure 6.

**Supplemental Table 1:** p values for DSBs in different stages of oogenesis of indicated strains compared to their controls

| Control | Strain | S1 N.C.<sup>2</sup> | S1 O<sup>3</sup> | S2 N.C.<sup>2</sup> | S2 O<sup>3</sup> | S3 N.C.<sup>2</sup> | S3 O<sup>3</sup> | S4 N.C.<sup>2</sup> | S4 O<sup>3</sup> |
| --- | --- | --- | --- | --- | --- | --- | --- | --- | --- |
| Wild Type | p53A-B- | 0.0163 | 0.0193 | 0.02 | 0.02 | 0.17 | 0.17 | 0.04 | 0.04 |
| Wild Type | p53A-B+ | 0.42 | 0.05 | 1 | 1 | 1 | 1 | 1 | 1 |
| Wild Type | p53A+B- | 0.0001 | 0.0001 | N.D | 0.0011 | 1 | 1 | 1 | 1 |
| Wild Type | okra<sup>-</sup>;p53A+B+ | 0.1192 | 0.0002 | 0.154 | 0.0013 | 0.05 | 0.06 | N.D | N.D |
| Wild Type | okra<sup>-</sup>;p53A-B- | 0.0071 | 0.0028 | 0.0003 | N.D | 0.0002 | 0.0008 | 0.0138 | 0.0005 |
| Wild Type | okra<sup>-</sup>;p53A-B+ | N.D | 0.0001 | N.D | N.D | N.D | 0.0028 | N.D | 0.0121 |
| Wild Type | okra<sup>-</sup>;p53A+B- | N.D | 0.0001 | N.D | N.D | N.D | N.D | N.D | N.D |
| okra<sup>-</sup> | okra<sup>-</sup>;p53A-B- | 0.9579 | 0.0054 | 0.0299 | 0.1295 | 0.0002 | 0.0185 | 0.0026 | 0.0001 |
| okra<sup>-</sup> | okra<sup>-</sup>;p53A-B+ | 0.7381 | 0.2366 | 0.154 | 0.2366 | 0.05 | 0.06 | N.D | 0.0012 |
| okra<sup>-</sup> | okra<sup>-</sup>;p53A+B- | 0.1959 | 0.2366 | 0.0653 | 0.2366 | 0.0005 | 0.0411 | N.D | N.D |

<sup>1</sup> Relevant to Figures 7 and 8, p values calculated using unpaired student's t test
<sup>2</sup>N.C: Nurse cells
<sup>3</sup>O: Oocyte
<sup>4</sup> p values in **red font** are significant and those in black are not significant
<sup>5</sup>N.D: Not Determined since SEM is zero

**Supplemental Table 2:**
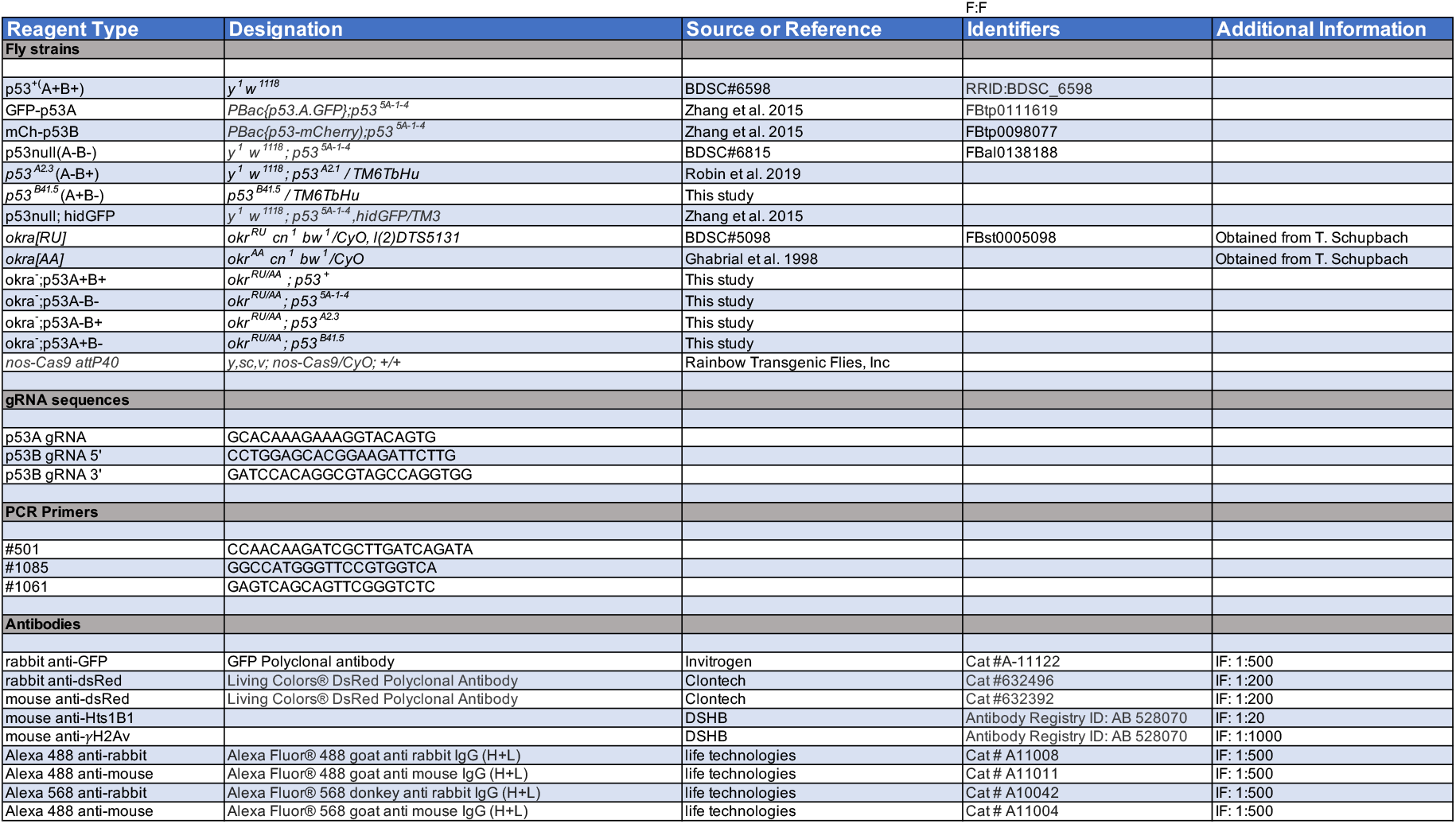
Key Resources.

## Notes

### Competing Interest Statement

The authors have declared no competing interest.

### Summary of Updates

This version corrects typographical errors discovered in the previous version.

## References

Alberti S. 2017. Phase separation in biology. Curr Biol 27: R1097–R1102.

Anbarasan T, Bourdon JC. 2019. The Emerging Landscape of p53 Isoforms in Physiology, Cancer and Degenerative Diseases. International journal of molecular sciences 20.

Aoubala M, Murray-Zmijewski F, Khoury MP, Fernandes K, Perrier S, Bernard H, Prats AC, Lane DP, Bourdon JC. 2011. p53 directly transactivates Delta133p53alpha, regulating cell fate outcome in response to DNA damage. Cell Death Differ 18: 248–258.

Bahler J, Hagens G, Holzinger G, Scherthan H, Heyer WD. 1994. Saccharomyces cerevisiae cells lacking the homologous pairing protein p175SEP1 arrest at pachytene during meiotic prophase. Chromosoma 103: 129–141.

Bolcun-Filas E, Rinaldi VD, White ME, Schimenti JC. 2014. Reversal of female infertility by Chk2 ablation reveals the oocyte DNA damage checkpoint pathway. Science 343: 533–536.

Brodsky MH, Nordstrom W, Tsang G, Kwan E, Rubin GM, Abrams JM. 2000. Drosophila p53 binds a damage response element at the reaper locus. Cell 101: 103–113.

Brodsky MH, Weinert BT, Tsang G, Rong YS, McGinnis NM, Golic KG, Rio DC, Rubin GM. 2004. Drosophila melanogaster MNK/Chk2 and p53 regulate multiple DNA repair and apoptotic pathways following DNA damage. Mol Cell Biol 24: 1219–1231.

Calvi BR, Lilly MA, Spradling AC. 1998. Cell cycle control of chorion gene amplification. Genes Dev 12: 734–744.

Candi E, Agostini M, Melino G, Bernassola F. 2014. How the TP53 family proteins TP63 and TP73 contribute to tumorigenesis: regulators and effectors. Human mutation 35: 702–714.

Carpenter AT. 1975. Electron microscopy of meiosis in Drosophila melanogaster females. I. Structure, arrangement, and temporal change of the synaptonemal complex in wild-type. Chromosoma 51: 157–182.

Chang HR, Munkhjargal A, Kim MJ, Park SY, Jung E, Ryu JH, Yang Y, Lim JS, Kim Y. 2018. The functional roles of PML nuclear bodies in genome maintenance. Mutation research 809: 99–107.

Coutandin D, Osterburg C, Srivastav RK, Sumyk M, Kehrloesser S, Gebel J, Tuppi M, Hannewald J, Schafer B, Salah E et al. 2016. Quality control in oocytes by p63 is based on a spring-loaded activation mechanism on the molecular and cellular level. eLife 5.

de la Cova C, Senoo-Matsuda N, Ziosi M, Wu DC, Bellosta P, Quinzii CM, Johnston LA. 2014. Supercompetitor status of Drosophila Myc cells requires p53 as a fitness sensor to reprogram metabolism and promote viability. Cell metabolism 19: 470–483.

Dej KJ, Spradling AC. 1999. The endocycle controls nurse cell polytene chromosome structure during Drosophila oogenesis. Development 126: 293–303.

Deng WM, Althauser C, Ruohola-Baker H. 2001. Notch-Delta signaling induces a transition from mitotic cell cycle to endocycle in Drosophila follicle cells. Development 128: 4737–4746.

Di Giacomo M, Barchi M, Baudat F, Edelmann W, Keeney S, Jasin M. 2005. Distinct DNA-damage-dependent and -independent responses drive the loss of oocytes in recombination-defective mouse mutants. Proc Natl Acad Sci U S A 102: 737–742.

Dichtel-Danjoy ML, Ma D, Dourlen P, Chatelain G, Napoletano F, Robin M, Corbet M, Levet C, Hafsi H, Hainaut P et al. 2013. Drosophila p53 isoforms differentially regulate apoptosis and apoptosis-induced proliferation. Cell Death Differ 20: 108–116.

Dotsch V, Bernassola F, Coutandin D, Candi E, Melino G. 2010. p63 and p73, the ancestors of p53. Cold Spring Harbor perspectives in biology 2: a004887.

Drummond-Barbosa D. 2019. Local and Physiological Control of Germline Stem Cell Lineages in Drosophila melanogaster. Genetics 213: 9–26.

Fogal V, Gostissa M, Sandy P, Zacchi P, Sternsdorf T, Jensen K, Pandolfi PP, Will H, Schneider C, Del Sal G. 2000. Regulation of p53 activity in nuclear bodies by a specific PML isoform. The EMBO journal 19: 6185–6195.

Fujita K. 2019. p53 Isoforms in Cellular Senescence- and Ageing-Associated Biological and Physiological Functions. International journal of molecular sciences 20.

Fujita K, Mondal AM, Horikawa I, Nguyen GH, Kumamoto K, Sohn JJ, Bowman ED, Mathe EA, Schetter AJ, Pine SR et al. 2009. p53 isoforms Delta133p53 and p53beta are endogenous regulators of replicative cellular senescence. Nature cell biology 11: 1135–1142.

Ge DT, Tipping C, Brodsky MH, Zamore PD. 2016. Rapid Screening for CRISPR-Directed Editing of the Drosophila Genome Using white Coconversion. G3 6: 3197–3206.

Gebel J, Tuppi M, Krauskopf K, Coutandin D, Pitzius S, Kehrloesser S, Osterburg C, Dotsch V. 2017. Control mechanisms in germ cells mediated by p53 family proteins. Journal of cell science.

Ghabrial A, Ray RP, Schupbach T. 1998. okra and spindle-B encode components of the RAD52 DNA repair pathway and affect meiosis and patterning in Drosophila oogenesis. Genes Dev 12: 2711–2723.

Ghabrial A, Schupbach T. 1999. Activation of a meiotic checkpoint regulates translation of Gurken during Drosophila oogenesis. Nature cell biology 1: 354–357.

Gong L, Gong H, Pan X, Chang C, Ou Z, Ye S, Yin L, Yang L, Tao T, Zhang Z et al. 2015. p53 isoform Delta113p53/Delta133p53 promotes DNA double-strand break repair to protect cell from death and senescence in response to DNA damage. Cell research 25: 351–369.

Gratz SJ, Wildonger J, Harrison MM, O’Connor-Giles KM. 2013. CRISPR/Cas9-mediated genome engineering and the promise of designer flies on demand. Fly 7.

Hassel C, Zhang B, Dixon M, Calvi BR. 2014. Induction of endocycles represses apoptosis independently of differentiation and predisposes cells to genome instability. Development 141: 112–123.

Hinnant TD, Merkle JA, Ables ET. 2020. Coordinating Proliferation, Polarity, and Cell Fate in the Drosophila Female Germline. Front Cell Dev Biol 8: 19.

Hughes SE, Miller DE, Miller AL, Hawley RS. 2018. Female Meiosis: Synapsis, Recombination, and Segregation in Drosophila melanogaster. Genetics 208: 875–908.

Ingaramo MC, Sanchez JA, Dekanty A. 2018. Regulation and function of p53: A perspective from Drosophila studies. Mechanisms of development 154: 82–90.

Jang JK, Sherizen DE, Bhagat R, Manheim EA, McKim KS. 2003. Relationship of DNA double-strand breaks to synapsis in Drosophila. Journal of cell science 116: 3069–3077.

Jia D, Huang YC, Deng WM. 2015. Analysis of Cell Cycle Switches in Drosophila Oogenesis. Methods Mol Biol 1328: 207–216.

Jin S, Martinek S, Joo WS, Wortman JR, Mirkovic N, Sali A, Yandell MD, Pavletich NP, Young MW, Levine AJ. 2000. Identification and characterization of a p53 homologue in Drosophila melanogaster. Proc Natl Acad Sci U S A 97: 7301–7306.

Joruiz SM, Bourdon JC. 2016. p53 Isoforms: Key Regulators of the Cell Fate Decision. Cold Spring Harbor perspectives in medicine.

Jost CA, Marin MC, Kaelin WG, Jr. 1997. p73 is a simian [correction of human] p53-related protein that can induce apoptosis. Nature 389: 191–194.

Joyce EF, McKim KS. 2009. Drosophila PCH2 is required for a pachytene checkpoint that monitors double-strand-break-independent events leading to meiotic crossover formation. Genetics 181: 39–51.

Joyce EF, McKim KS. 2011. Meiotic checkpoints and the interchromosomal effect on crossing over in Drosophila females. Fly 5: 134–140.

King RC. 1970. Ovarian Development in Drosophila melanogaster. Academic Press, New York.

Lake CM, Korda Holsclaw J, Bellendir SP, Sekelsky J, Hawley RS. 2013. The Development of a Monoclonal Antibody Recognizing the Drosophila melanogaster Phosphorylated Histone H2A Variant (gamma-H2AV). G3.

Lane DP, Crawford LV. 1979. T antigen is bound to a host protein in SV40-transformed cells. Nature 278: 261–263.

Levine AJ. 2019. The many faces of p53: something for everyone. J Mol Cell Biol 11: 524–530.

Levine AJ. 2020. p53: 800 million years of evolution and 40 years of discovery. Nature reviews Cancer.

Li XC, Schimenti JC. 2007. Mouse pachytene checkpoint 2 (trip13) is required for completing meiotic recombination but not synapsis. PLoS Genet 3: e130.

Lin H, Spradling AC. 1993. Germline stem cell division and egg chamber development in transplanted Drosophila germaria. Dev Biol 159: 140–152.

Lin H, Yue L, Spradling A. 1994. The Drosophila fusome, a germline-specific organelle, contains membrane skeletal proteins and functions in cyst formation. Development 120: 947–956.

Linke SP, Sengupta S, Khabie N, Jeffries BA, Buchhop S, Miska S, Henning W, Pedeux R, Wang XW, Hofseth LJ et al. 2003. p53 interacts with hRAD51 and hRAD54, and directly modulates homologous recombination. Cancer Res 63: 2596–2605.

Linzer DI, Levine AJ. 1979. Characterization of a 54K dalton cellular SV40 tumor antigen present in SV40-transformed cells and uninfected embryonal carcinoma cells. Cell 17: 43–52.

Liu JL, Buszczak M, Gall JG. 2006. Nuclear bodies in the Drosophila germinal vesicle. Chromosome research: an international journal on the molecular, supramolecular and evolutionary aspects of chromosome biology 14: 465–475.

Liu JL, Wu Z, Nizami Z, Deryusheva S, Rajendra TK, Beumer KJ, Gao H, Matera AG, Carroll D, Gall JG. 2009. Coilin is essential for Cajal body organization in Drosophila melanogaster. Molecular biology of the cell 20: 1661–1670.

Lu WJ, Chapo J, Roig I, Abrams JM. 2010. Meiotic recombination provokes functional activation of the p53 regulatory network. Science 328: 1278–1281.

Ma X, Han Y, Song X, Do T, Yang Z, Ni J, Xie T. 2016. DNA damage-induced Lok/CHK2 activation compromises germline stem cell self-renewal and lineage differentiation. Development 143: 4312–4323.

Madigan JP, Chotkowski HL, Glaser RL. 2002. DNA double-strand break-induced phosphorylation of Drosophila histone variant H2Av helps prevent radiation-induced apoptosis. Nucleic Acids Res 30: 3698–3705.

Marcet-Ortega M, Pacheco S, Martinez-Marchal A, Castillo H, Flores E, Jasin M, Keeney S, Roig I. 2017. p53 and TAp63 participate in the recombination-dependent pachytene arrest in mouse spermatocytes. PLoS Genet 13: e1006845.

Mateo AR, Kessler Z, Jolliffe AK, McGovern O, Yu B, Nicolucci A, Yanowitz JL, Derry WB. 2016. The p53-like Protein CEP-1 Is Required for Meiotic Fidelity in C. elegans. Curr Biol 26: 1148–1158.

Mauri F, McNamee LM, Lunardi A, Chiacchiera F, Del Sal G, Brodsky MH, Collavin L. 2008. Modification of Drosophila p53 by SUMO modulates its transactivation and pro-apoptotic functions. J Biol Chem 283: 20848–20856.

Mehrotra S, Maqbool SB, Kolpakas A, Murnen K, Calvi BR. 2008. Endocycling cells do not apoptose in response to DNA rereplication genotoxic stress. Genes Dev 22: 3158–3171.

Mehrotra S, McKim KS. 2006. Temporal analysis of meiotic DNA double-strand break formation and repair in Drosophila females. PLoS Genet 2: e200.

Mitrea DM, Kriwacki RW. 2016. Phase separation in biology; functional organization of a higher order. Cell Commun Signal 14: 1.

Monk AC, Abud HE, Hime GR. 2012. Dmp53 is sequestered to nuclear bodies in spermatogonia of Drosophila melanogaster. Cell and tissue research 350: 385–394.

Morris LX, Spradling AC. 2011. Long-term live imaging provides new insight into stem cell regulation and germline-soma coordination in the Drosophila ovary. Development 138: 2207–2215.

Moureau S, Luessing J, Harte EC, Voisin M, Lowndes NF. 2016. A role for the p53 tumour suppressor in regulating the balance between homologous recombination and non-homologous end joining. Open Biol 6.

Napoletano F, Gibert B, Yacobi-Sharon K, Vincent S, Favrot C, Mehlen P, Girard V, Teil M, Chatelain G, Walter L et al. 2017. p53-dependent programmed necrosis controls germ cell homeostasis during spermatogenesis. PLoS Genet 13: e1007024.

Ollmann M, Young LM, Di Como CJ, Karim F, Belvin M, Robertson S, Whittaker K, Demsky M, Fisher WW, Buchman A et al. 2000. Drosophila p53 is a structural and functional homolog of the tumor suppressor p53. Cell 101: 91–101.

Page SL, Hawley RS. 2001. c(3)G encodes a Drosophila synaptonemal complex protein. Genes Dev 15: 3130–3143.

Park JH, Nguyen TTN, Lee EM, Castro-Aceituno V, Wagle R, Lee KS, Choi J, Song YH. 2019. Role of p53 isoforms in the DNA damage response during Drosophila oogenesis. Scientific reports 9: 11473.

Qi S, Calvi BR. 2016. Different cell cycle modifications repress apoptosis at different steps independent of developmental signaling in Drosophila. Molecular biology of the cell 27: 1885–1897.

Ren X, Sun J, Housden BE, Hu Y, Roesel C, Lin S, Liu LP, Yang Z, Mao D, Sun L et al. 2013. Optimized gene editing technology for Drosophila melanogaster using germ line-specific Cas9. Proc Natl Acad Sci U S A 110: 19012–19017.

Rinaldi VD, Bloom JC, Schimenti JC. 2020. Oocyte Elimination Through DNA Damage Signaling from CHK1/CHK2 to p53 and p63. Genetics 215: 373–378.

Rinaldi VD, Bolcun-Filas E, Kogo H, Kurahashi H, Schimenti JC. 2017. The DNA Damage Checkpoint Eliminates Mouse Oocytes with Chromosome Synapsis Failure. Mol Cell 67: 1026–1036 e1022.

Robin M, Issa AR, Santos CC, Napoletano F, Petitgas C, Chatelain G, Ruby M, Walter L, Birman S, Domingos PM et al. 2019. Drosophila p53 integrates the antagonism between autophagy and apoptosis in response to stress. Autophagy 15: 771–784.

Roy S, Ernst J, Kharchenko PV, Kheradpour P, Negre N, Eaton ML, Landolin JM, Bristow CA, Ma L, Lin MF et al. 2010. Identification of functional elements and regulatory circuits by Drosophila modENCODE. Science 330: 1787–1797.

Rutkowski R, Hofmann K, Gartner A. 2010. Phylogeny and function of the invertebrate p53 superfamily. Cold Spring Harbor perspectives in biology 2: a001131.

San-Segundo PA, Roeder GS. 1999. Pch2 links chromatin silencing to meiotic checkpoint control. Cell 97: 313–324.

Sekelsky J. 2017. DNA Repair in Drosophila: Mutagens, Models, and Missing Genes. Genetics 205: 471–490.

Sogame N, Kim M, Abrams JM. 2003. Drosophila p53 preserves genomic stability by regulating cell death. Proc Natl Acad Sci U S A 100: 4696–4701.

Spradling AC. 1993. Developmental Genetics of Oogenesis. in The Development of Drosophila melanogaster (ed. MBaA Martinez-Arias). Cold Spring Harbor Press, Cold Spring Harbor, New York.

Suh EK, Yang A, Kettenbach A, Bamberger C, Michaelis AH, Zhu Z, Elvin JA, Bronson RT, Crum CP, McKeon F. 2006. p63 protects the female germ line during meiotic arrest. Nature 444: 624–628.

Tanaka-Matakatsu M, Xu J, Cheng L, Du W. 2009. Regulation of apoptosis of rbf mutant cells during Drosophila development. Dev Biol 326: 347–356.

Tasnim S, Kelleher ES. 2018. p53 is required for female germline stem cell maintenance in P-element hybrid dysgenesis. Dev Biol 434: 215–220.

Thomer M, May NR, Aggarwal BD, Kwok G, Calvi BR. 2004. Drosophila double-parked is sufficient to induce re-replication during development and is regulated by Cyclin E / CDK2. Development 131: 4807–4818.

Thurmond J, Goodman JL, Strelets VB, Attrill H, Gramates LS, Marygold SJ, Matthews BB, Millburn G, Antonazzo G, Trovisco V et al. 2019. FlyBase 2.0: the next generation. Nucleic Acids Res 47: D759–D765.

Wei Y, Bettedi L, Ting CY, Kim K, Zhang Y, Cai J, Lilly MA. 2019. The GATOR complex regulates an essential response to meiotic double-stranded breaks in Drosophila. eLife 8.

Wells BS, Yoshida E, Johnston LA. 2006. Compensatory proliferation in Drosophila imaginal discs requires Dronc-dependent p53 activity. Curr Biol 16: 1606–1615.

White AE, Leslie ME, Calvi BR, Marzluff WF, Duronio RJ. 2007. Developmental and cell cycle regulation of the Drosophila histone locus body. Molecular biology of the cell 18: 2491–2502.

Williams AB, Schumacher B. 2016. p53 in the DNA-Damage-Repair Process. Cold Spring Harbor perspectives in medicine 6.

Wu HY, Burgess SM. 2006. Two distinct surveillance mechanisms monitor meiotic chromosome metabolism in budding yeast. Curr Biol 16: 2473–2479.

Wylie A, Jones AE, D’Brot A, Lu WJ, Kurtz P, Moran JV, Rakheja D, Chen KS, Hammer RE, Comerford SA et al. 2016. p53 genes function to restrain mobile elements. Genes Dev 30: 64–77.

Wylie A, Lu WJ, D’Brot A, Buszczak M, Abrams JM. 2014. p53 activity is selectively licensed in the Drosophila stem cell compartment. eLife 3: e01530.

Xing Y, Su TT, Ruohola-Baker H. 2015. Tie-mediated signal from apoptotic cells protects stem cells in Drosophila melanogaster. Nature communications 6: 7058.

Yang A, Kaghad M, Wang Y, Gillett E, Fleming MD, Dotsch V, Andrews NC, Caput D, McKeon F. 1998. p63, a p53 homolog at 3q27-29, encodes multiple products with transactivating, death-inducing, and dominant-negative activities. Mol Cell 2: 305–316.

Zaccai M, Lipshitz HD. 1996. Differential distributions of two adducin-like protein isoforms in the Drosophila ovary and early embryo. Zygote 4: 159–166.

Zhang B, Mehrotra S, Ng WL, Calvi BR. 2014. Low Levels of p53 Protein and Chromatin Silencing of p53 Target Genes Repress Apoptosis in Drosophila Endocycling Cells. PLoS Genet 10: e1004581.

Zhang B, Rotelli M, Dixon M, Calvi BR. 2015. The function of Drosophila p53 isoforms in apoptosis. Cell Death Differ 22: 2058–2067.

Zhou L. 2019. P53 and Apoptosis in the Drosophila Model. Advances in experimental medicine and biology 1167: 105–112.

